# Shift of lung macrophage composition is associated with COVID-19 disease severity and recovery

**DOI:** 10.1101/2022.01.11.475918

**Authors:** Steven T. Chen, Matthew D. Park, Diane Marie Del Valle, Mark Buckup, Alexandra Tabachnikova, Nicole W. Simons, Konstantinos Mouskas, Brian Lee, Daniel Geanon, Darwin D’Souza, Travis Dawson, Robert Marvin, Kai Nie, Ryan C. Thompson, Zhen Zhao, Jessica LeBerichel, Christie Chang, Hajra Jamal, Udit Chaddha, Kusum Mathews, Samuel Acquah, Stacey-Ann Brown, Michelle Reiss, Timothy Harkin, Marc Feldmann, Charles A. Powell, Jaime L. Hook, Seunghee Kim-Schulze, Adeeb H. Rahman, Brian D. Brown, The Mount Sinai COVID-19 Biobank Team, Noam D. Beckmann, Sacha Gnjatic, Ephraim Kenigsberg, Alexander W. Charney, Miriam Merad

**Author notes:** Indicates equal contribution (Sacha Gnjatic, Ephraim Kenigsberg, Alexander W. Charney, Miriam Merad).

## Abstract

Though it has been 2 years since the start of the Coronavirus Disease 19 (COVID-19) pandemic, COVID-19 continues to be a worldwide health crisis. Despite the development of preventive vaccines, very little progress has been made to identify curative therapies to treat COVID-19 and other inflammatory diseases which remain a major unmet need in medicine. Our study sought to identify drivers of disease severity and death to develop tailored immunotherapy strategies to halt disease progression. Here we assembled the Mount Sinai COVID-19 Biobank which was comprised of ~600 hospitalized patients followed longitudinally during the peak of the pandemic. Moderate disease and survival were associated with a stronger antigen (Ag) presentation and effector T cell signature, while severe disease and death were associated with an altered Ag presentation signature, increased numbers of circulating inflammatory, immature myeloid cells, and extrafollicular activated B cells associated with autoantibody formation. Strikingly, we found that in severe COVID-19 patients, lung tissue resident alveolar macrophages (AM) were not only severely depleted, but also had an altered Ag presentation signature, and were replaced by inflammatory monocytes and monocyte-derived macrophages (MoMΦ). Notably, the size of the AM pool correlated with recovery or death, while AM loss and functionality were restored in patients that recovered. These data therefore suggest that local and systemic myeloid cell dysregulation is a driver of COVID-19 severity and that modulation of AM numbers and functionality in the lung may be a viable therapeutic strategy for the treatment of critical lung inflammatory illnesses.

## INTRODUCTION

Though there has been unprecedented success with the concurrent development of multiple highly effective preventive vaccines against SARS-CoV-2, there remains a critical need to develop novel, targeted immune therapies for vulnerable populations and critically ill patients as vaccine rollout continues worldwide and potential breakthrough variants continue to arise. In addition to COVID-19, there is a critical need to develop novel therapies to recognize, modulate, and treat pathogenic inflammation associated with critical inflammatory illnesses, especially in the elderly population.

Here we assembled the Mount Sinai COVID-19 Biobank, which collected longitudinal blood and tissue samples from patients and healthy controls, and investigated local immune dynamics in the lungs of infected patients *(1)*. 583 COVID-19 patients, who were hospitalized at the Mount Sinai Hospital, were enrolled into the Mount Sinai COVID-19 Biobank (**Table S1**). Serum and peripheral blood mononuclear cells (PBMC) were collected from patients on timepoint 1 (T1), on average 14.8±10.6 days self-reported post-symptom onset (PSO). Samples were assigned timepoint numbers according to approximately how many days post-hospitalization the sample was collected (e.g. 4 days after hospitalization= T4). Severely ill patients, who were hospitalized >2 weeks, had an additional sample collected 7 days later (T13). Severity scoring for each patient sample was assigned using clinical criteria designated by Mount Sinai Hospital (**Table S2**)

## RESULTS

### Proteomic characterization of COVID-19 sera reveals distinct immune patterns

To characterize the diversity of immune patterns in COVID-19 patients, we measured 92 different cytokines on 1956 COVID^+^ and 45 healthy donor (HD) COVID^-^ serum samples using the Olink inflammation panel. Instead of solely relying on clinical severity to group patients, we performed unsupervised clustering to unbiasedly sort sera into 15 different cytokine patterns (**Fig. 1A-C, Fig. S1A-B**). The majority of patients had 1-4 timepoints which were distributed across all cytokine patterns (**Fig. S2A-D**). Strikingly, we found that the immune patterns were associated with clinical severity and final patient outcome, leading us to group them. Group 1 consisted of patterns 12-15 and was enriched in samples from HD and Moderate COVID-19 patients; Group 2, which included patterns 6-9, was our largest and most heterogeneous group, but was enriched in Severe COVID-19 samples. Immune patterns 8-9 had increased levels of Interferon γ (IFNγ) responsive and T helper type 1 (Th1) activation cytokines [e.g. IFNγ, C-X-C Motif Chemokine ligand 9, 10, 11 (CXCL9, CXCL10, CXCL11), IL-2)] compared to clusters 6-7*(2)*. Group 3, patterns 1-5, was enriched in Severe COVID-19 with end organ damage (EOD) samples, as well as samples from patients that died.

**Figure 1.**
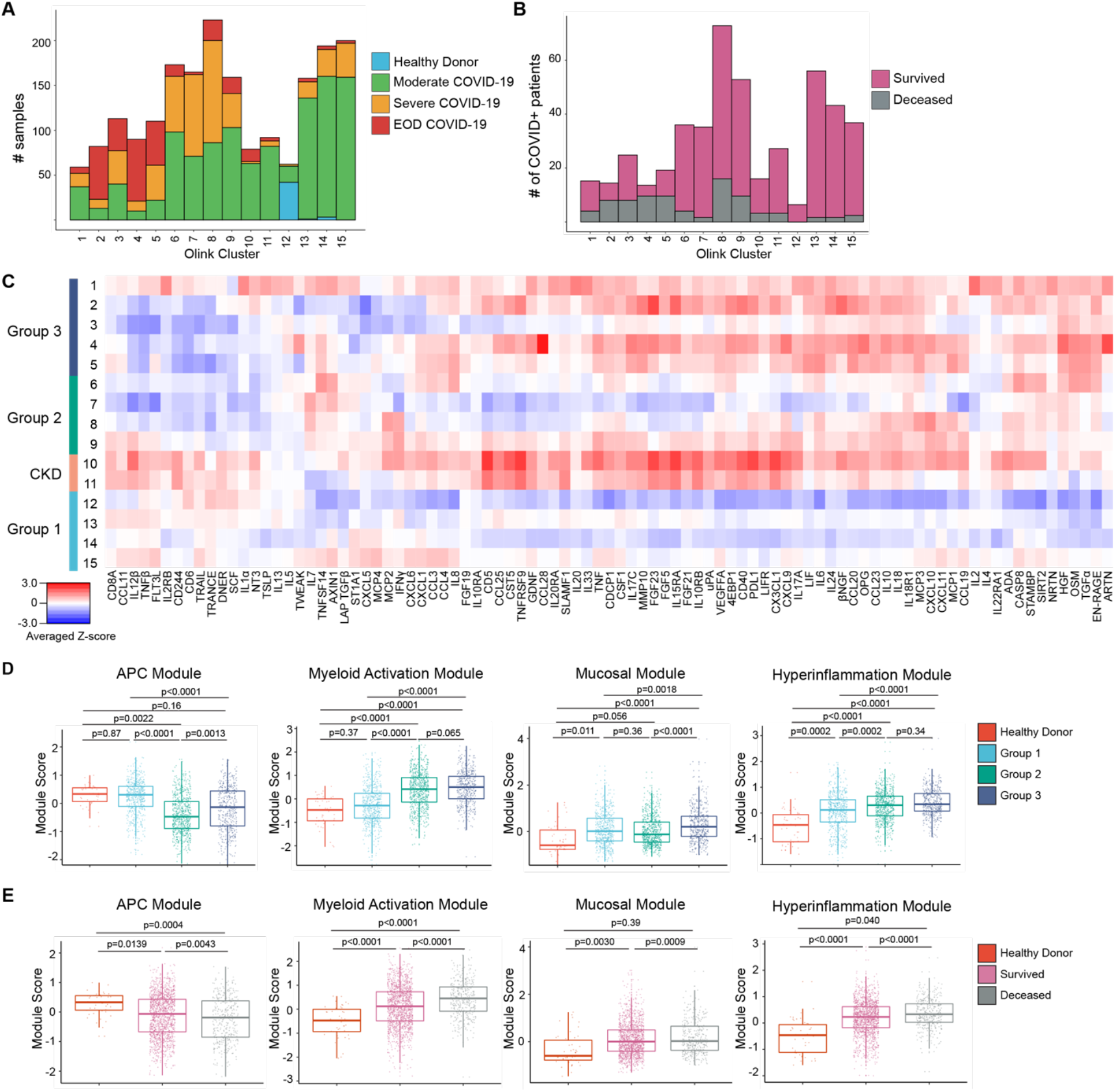
Proteomic characterization of COVID-19 serum reveals distinct immune patterns associated with disease severity and clinical outcome. **(A)** Histogram of patient samples across Olink clusters denoted by clinical severity classification. (**B)** Histogram of first available patient samples across Olink clusters denoted by patient projected clinical outcome. (**C)** Averaged z-scored heatmap of Olink inflammation panel analytes across Olink clusters. Olink clusters were grouped based on clinical severity, projected outcome, and comorbidity distribution. (**D)** Boxplots showing Olink module score comparisons of all Olink samples by Olink group. (**E)** Boxplots showing Olink module score comparisons of all Olink samples by final clinical outcome. For box plots, each dot represents a patient sample; center line, median; box limits, 25^th^ and 75^th^ percentile; whiskers, 1.5x interquartile range (IQR). Statistical significance (**D-E**) determined by 2-way ANOVA with Tukey’s Multiple Comparisons correction. Adjusted p-values shown.

Our unbiased clustering was not driven by sex, body mass index (BMI), or smoking status, but age and days PSO at time of sampling were higher in Group 3 samples (**Fig. S2E-I**). Group 2 (patterns 6-9) and 3 (patterns 1-5) patients had higher concentrations of C-reactive protein and D-Dimer, indicating increased inflammation and hypercoagulability (**Fig. S2J-K**). Hypertension (HTN) and Diabetes (DM) were common comorbidities within our cohort, especially in Group 3 (**Fig. S2L**). Immune patterns 10-11 were highly enriched in patients with chronic kidney disease (CKD), HTN, DM, and heart failure, leading us to group them into a distinct CKD group. Almost all patients within our cohort received anticoagulation, and patients in Group 2 and 3 were more likely to receive steroids (**Fig. S2M**). Notably, while this unbiased clustering showed that immune patterns could be grouped based on enrichment of samples from clinical severity classification, patient samples with similar clinical parameters were assigned to diverse cytokine-based patterns, indicating that clinical scoring was unable to fully capture the diversity of immune patterns in COVID-19.

Based on the covariance patterns of the Olink cytokines (**Fig. S3**), we identified 4 protein modules and calculated module scores for each Olink group. The antigen presenting cell (APC) module, which included proteins associated with Ag presentation, dendritic cells (DC), and T cell activation [Tumor necrosis factor (TNF)-related apoptosis-inducing ligand (TRAIL), TNF-related activation-induced cytokine (TRANCE), IL-12β, FMS-like tyrosine kinase 3 ligand (FLT3L), TNF beta (TNFβ)], scored higher in patients with moderate disease and healthy controls (Fig. 1D-E)*(3–6)*. Next, we identified a core group of 4 cytokines released by activated monocytes and neutrophils, [Transforming growth factor alpha (TGFα), Hepatocyte growth factor (HGF), Oncostatin M (OSM), S100 calcium-binding protein A12 (EN-RAGE/S100A12)] which were enriched in patients with Severe or EOD COVID-19 and grouped them into a myeloid activation module. Signaling by these cytokines have been associated with proinflammatory cytokine secretion, fibroblast activation, and fibrosis*(7–19)*. The mucosal module, which included T helper 17 (Th17) and barrier defense cytokines [IL-17A, IL-17C, C-C Motif Chemokine Ligand 20, 28 (CCL20, CCL28), IL-33], and the hyperinflammation module which included inflammatory cytokines [TNF, IL-6, IL-8, IL-10, IL-18, CXCL10, Monocyte chemoattractant protein-3 (MCP-3)] were more enriched in patients with severe or EOD disease*(20–22)*. We grouped these analytes into a Mucosal Module and a Hyperinflammation Module, respectively.

APC module scores were higher in healthy controls and Group 1 and reduced in Group 2 and 3, while Myeloid Activation, Mucosal, and Hyperinflammation module scores were higher in Groups 2 and 3(**Fig. 1D**). Comparison of module scores by final clinical outcome showed that patients who survived had a higher APC module score, while patients that died had higher Myeloid Activation, Mucosal, and Hyperinflammation module scores (**Fig. 1E**). These trends held even when we compared only the first timepoints for each patient, suggesting that module scores could be used early on during a patient’s hospitalization to predict clinical outcome (**Fig S4A-B**).

To determine the stability of these immune patterns, we performed a discrete time Markov chain analysis to measure the probability of transition between cytokine patterns across successive samples, irrespective of past or future states (**Fig. S4C**)*(23)*. Between timepoints, Group 1 patients had a higher probability of transitioning to other Group 1 immune patterns and a higher probability of survival compared to other groups. Group 2 patients also had a high probability of transitioning to Group 1 patterns 13-15. However, compared to Group 1 patients, Group 2 patients had a higher probability of death between timepoints. CKD patients had ~50% probability of survival or death between timepoints and a low probability of transitioning to either Group 1 or Group 2 immune patterns. Finally Group 3 patients had a high probability of remaining within Group 3 immune patterns between timepoints and the highest probability of dying.

This proteomic analysis highlights the heterogeneity of immune states in COVID-19 that remained stable over time, despite medical intervention, and the potential value of using Olink module scores to predict outcome and response to treatment. The heterogeneity revealed by our clustering underscores the limitations of solely using clinical severity parameters to stratify patients for treatment. For example, while CKD patients and Group 3 patients may benefit from broad immune suppression and targeted therapies like IL-6 blockade, these same treatments are unlikely to show the same effect in patients with Group 2 cytokine patterns, and may instead hinder protective adaptive immune responses. On the other hand, all patients would likely benefit from therapies to boost their APC response, such as administration of Flt3L, to increase the number of DC for T cell priming and activation.

### Myeloid Cell Dysregulation underlies COVID-19 Severity

We performed cytometry by time of flight (CyTOF) on whole blood samples to measure circulating immune cell composition and its association with Olink group immune patterns, Consistent with prior studies, we found that neutrophils, classical monocytes, and intermediate monocytes were significantly increased while all DC populations trended down in more severe disease. (**Fig. 2A-B**). Grouping patient samples by final clinical outcome showed that patients who died from COVID-19 had increased numbers of neutrophils (**Fig. 2C**). Classical and Intermediate monocytes were significantly increased in all COVID-19 patients while DC populations trended down relative to HD (**Fig. 2D**). CyTOF also confirmed lymphopenia of both CD4 and CD8 T cells in Group 2 and 3 patients (**Fig. S5A-B**)*(24, 25)*. Naïve, central memory (CM), and effector memory (EM) and effector memory re-expressing CD45RA (EMRA) CD4 and CD8 T cells also trended downwards in more severely ill COVID-19 patients.

**Figure 2.**
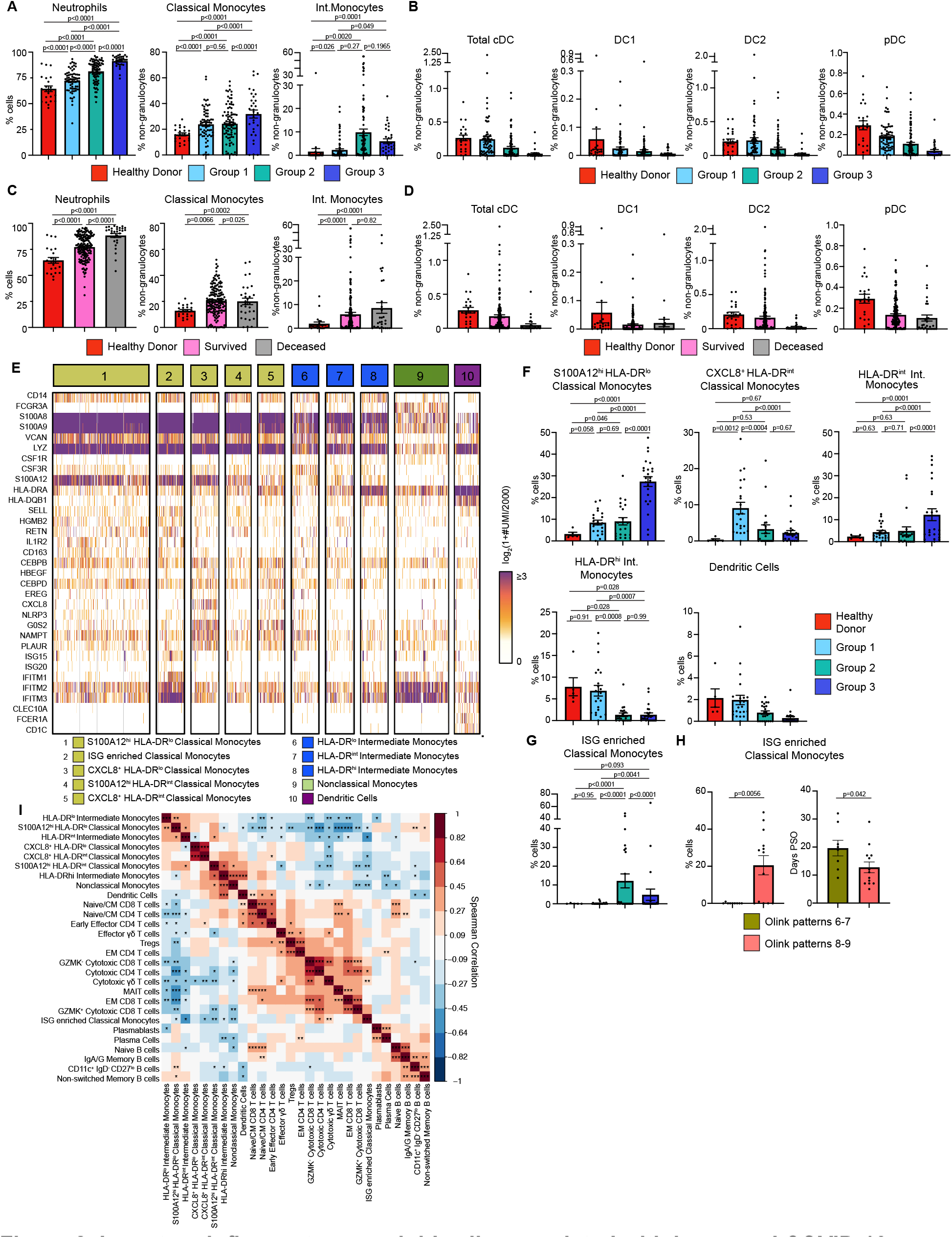
Immature inflammatory myeloid cells associated with increased COVID-19 severity. **(A)** Neutrophils (% cells), Classical Monocytes and Intermediate Monocyte frequencies (% non-granulocytes) in whole blood by Olink group measured by CyTOF. **(B)** DC population frequencies (% non-granulocytes) in whole blood by Olink group measured by CyTOF. **(C)** Neutrophils (% cells), Classical Monocytes and Intermediate Monocyte frequencies (%non-granulocytes) in whole blood by final clinical outcome measured by CyTOF. **(D)** DC population frequencies (% non-granulocytes) in whole blood by final clinical outcome measured by CyTOF. Conventional DC (cDC), conventional type 1 DC (DC1), conventional type 2 DC (DC2), and plasmacytoid DC (pDC) shown. **(E)** Heatmap showing unique molecular identifier (UMI) counts of selected genes from myeloid cell scRNAseq clusters from PBMC. **(F)** scRNAseq cluster cell frequencies as % of cells by Olink group **(G)** % of cells frequencies of ISG enriched Classical Monocytes cluster by Olink group. **(H)** % of cells frequencies and days PSO for ISG enriched Classical Monocytes by clusters 6-7 vs clusters 8-9. **(I)** Matrix of spearman correlation coefficients between identified scRNAseq PBMC clusters. *p<0.05, **p<0.005, ***p<0.0005. For bar graphs, each dot (**A-D, F-H**) represents a patient sample. Statistical significance (**A-D, F-G**) determined by 2-way ANOVA with Holm-Sidak multiple comparisons correction. Adjusted p-values shown. Statistical significance (**H**) determined by Mann-Whitney test.

To unbiasedly dissect the heterogeneity of immune cells, we performed single cell RNA sequencing (scRNAseq) on 81 PBMC samples from 39 COVID-19 patients and 6 healthy controls. After down-sampling, integration and batch correction, and removal of doublet cells, unsupervised clustering revealed discrete subsets of mononuclear phagocytes (MNP), T cells, and B cells (**Fig. 2E**). We identified a cluster of classical monocytes that highly expressed *S100A12* (EN-RAGE), *S100A8*, and *S100A9*, but lowly expressed human leukocyte antigen (HLA) molecules. Low expression of HLA molecules, *CSF1R* and concurrent high expression of granulocyte/monocyte precursor genes (i.e. *CSF3R, CEBPB, CEBPD*) suggested that this cluster was a group of immature cells arising from granulocyte-monocyte progenitors (GMP)*(26–29)*. This cluster, which we named S100A12^hi^ HLA-DR^lo^ Classical Monocytes, was found at significantly higher levels in Group 3 patients (**Fig. 2F**). Using this criteria for immature myeloid cells, we next identified 3 clusters of inflammatory immature monocytes expressing high levels of *S100A12*, inflammasome protein *NLRP3*, and oxidative stress marker *NAMPT(30, 31)*. These 3 monocyte clusters also expressed high levels of urokinase receptor, *PLAUR*, a marker of inflammation and fibrinolysis*(32)*. We distinguished these 3 clusters by levels of C-X-C Chemokine Motif Ligand 8 (CXCL8) and HLA-DR expression, naming them CXCL8^+^ HLA-DR^lo^ Classical Monocytes, S100A12^hi^ HLA-DR^int^ Classical Monocytes and CXCL8^+^ HLA-DR^int^ Classical Monocytes. CXCL8^+^ HLA-DR^int^ Classical Monocytes were specific to COVID-19 patients but were also found at higher levels in Group 1 patients compared to Group 2 and 3 patients. Taken together, our data supports previous work that immature, inflammatory myeloid cells that are likely arising from emergency myelopoiesis, is associated with increased COVID-19 severity*(27, 28)*.

We also identified a cluster of classical monocytes expressing high levels of type I IFN stimulated genes (ISG) (i.e. *ISG15, ISG20, IFITM1-3)* which was found at significantly higher numbers in Group 2 patients compared to Group.1 and 3 (**Fig. 2G**). Given the differences we saw in T cell activation cytokines by Olink, we stratified this cluster of ISG-enriched Classical monocytes by Olink patterns 6-7 vs 8-9. This group of monocytes was only found in patterns 8-9, but this could due to transient or delayed IFN signaling captured by earlier sampling of these patients days PSO (**Fig. 2H**)*(29, 33)*.

Next, we performed unbiased clustering to identify T cell clusters. Naïve/CM CD4 and CD8 T cells expressed *CCR7, IL7R, LDHB, LTB, LEF1* and *TCF7* and were found at higher levels in healthy controls (**Fig. S5C-E**)*(34, 35)*. We identified a cluster of Early Effector CD4 T cells that expressed low levels of *KLRB1, CCL5*, and *GZMM*, a cluster of T regulatory cells (Tregs), and mucosal associated invariant T (MAIT) cells that expressed high levels of *KLRB1, NKG7, GZMK, GZMA*, and *CCL5*. Effector memory (EM) CD4 and CD8 T cells expressed intermediate levels of *IL7R* and *LTB* and low levels of *LEF1, CCR7, TCF7 (34, 35)*. We also identified GZMK^+^, GZMK^-^ Cytotoxic CD8 T cells, and Cytotoxic CD4 T cells, as well as Cytotoxic and Effector γδ T cells based on granzyme and *GNLY* expression. Among circulating B cells, we identified Plasmablasts and Plasma cells by high *CD38, CD27, MZB1* expression; plasma cells were further distinguished by increased *PRDM1* expression (**Fig. S5F**) *(36, 37)*. *IGHD*^+^ and *IGHM*^+^ Naïve B cells were decreased in all COVID-19 patients. We also identified CD11c^+^ IgD^-^ CD27^lo^ B cells which are thought to be extrafollicular or polyreactive B cells that produce pathogenic autoantibodies (auto-Abs) (**Fig. S5G**)*(36, 38–40)*.

### Integration of circulating Immune cell phenotypes and serum proteomics

To probe how these different immune cell populations might be interacting, we performed a Spearman correlation analysis on the scRNAseq cell frequencies. Strikingly, we found that the frequencies of S100A12^hi^ HLA-DR^lo^ classical monocytes and other HLA-DR^lo^ immature monocyte clusters were negatively correlated with the ISG-enriched Classical Monocytes, DC, and cytotoxic T cell clusters, but were positively correlated with CD11 c^+^ IgD^-^ CD27^lo^ B cells (**Fig. 2I**). In contrast, DC were positively correlated with Naïve/CM CD4, CD8, Early Effector CD4 T cells, and Effector γδ T cells.

Next, we correlated scRNAseq cell composition and Olink proteomic profiles and hierarchically clustered cell subpopulations to group cell types with similar cytokine correlations (**Fig. 3**). Immature HLA-DR^lo^ myeloid cells, CD11c^+^ IgD^-^ CD27^lo^ B cells, and Non-switched Memory B cell frequencies were positively correlated with circulating levels of Myeloid Activation, Mucosal, and Hyperinflammatory module cytokines, and strongly negatively correlated with APC module cytokines. This immune program corresponded most closely with Group 3 patients who had increased numbers of circulating immature myeloid cells and higher concentrations of inflammatory cytokines. In contrast, DC clustered together with Naïve/CM CD4 and CD8 T cells, GZMK^+^ Cytotoxic CD8 T cells, Cytotoxic γδ T cells, MAIT cells, and Naïve B cells. This group of cell populations were positively correlated with APC module cytokines and negatively correlated with Myeloid Activation, Mucosal, and Hyperinflammation Module Cytokines. This pattern most closely corresponded to Group 1 patients who had higher levels of APC-T cell activation cytokines and effector T-cell populations.

**Figure 3:**
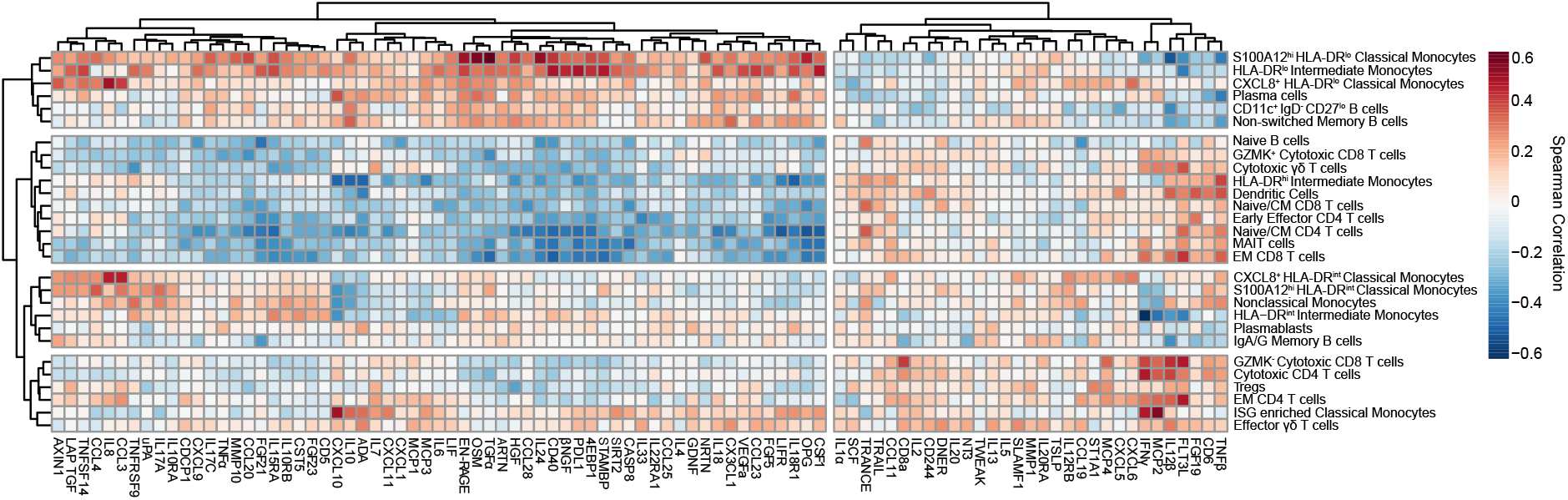
Integrated analysis of scRNAseq cluster frequencies and Olink analyte abundance in serum reveals distinct immune responses to COVID-19. Matrix of spearman correlation coefficients between identified scRNAseq PBMC clusters and Olink analyte normalized concentrations in serum. Axes ordered by hierarchical clustering.

We also identified two other patterns of cell populations which clustered together and were correlated with distinct cytokine profiles. HLA-DR’^nt^ monocyte populations clustered together with Nonclassical Monocytes, Plasmablasts, and IgA/IgG memory B cells. These cell types were negatively correlated with IFNγ, IL-12β and weakly negatively correlated with CXCL10 and CXCL11. In contrast, ISG-enriched Classical Monocytes clustered with EM CD4 T cells, GZMK^-^ Cytotoxic CD8 T cells, and Effector γδ T cells. These cell populations were positively correlated with IFNγ, IL-12β, CD8a, IL-2, and Flt3L concentrations, and negatively correlated with Myeloid Activation and Mucosal Module cytokines.

This integrated analysis showed 4 distinct types of immune response to COVID-19. First, Group 1 patients, who had higher numbers of more mature, HLA-DR^hi^ myeloid cell populations had correspondingly higher numbers of effector and cytotoxic T-cell populations, higher serum concentrations of APC and T cell activating cytokines, and reduced levels of inflammatory, tissue-damaging cytokines. These patients tended to have a milder course of COVID-19 and were more likely to recover. On the other end of the spectrum, Group 3 patients, who had high numbers of immature, HLA-DR^lo^ myeloid cells with limited Ag presentation capability, were likely unable to mount a strong T-cell response, and were instead more reliant on humoral control of infection. Immature myeloid cells in these patients may have predominated due to high levels of inflammatory cytokines that drive emergency myelopoiesis*(41, 42)*. These immature myeloid cells may have further contributed to hyperinflammation by the production of tissue-damaging cytokines and reactive oxygen species (ROS), leading to a vicious cycle of lymphopenia, a suppressed or delayed adaptive immune response, poor control of virus infection, and increased inflammation*(41, 42)*. Inflammation in these patients may also have been exacerbated by extrafollicular CD11c^+^ IgD^-^ CD27^lo^ B-cells secretion of auto-Abs that activated autoimmune inflammation*(39, 43, 44)*. Consequently, these patients had the lowest rates of survival. Patients with an earlier type I IFN response, as well as those who had higher numbers of mature HLA-DR^int/hi^ Monocytes and DC, may have been better protected against COVID-19 disease progression and morbidity because they were able to mount an earlier, more productive adaptive T cell response.

### Loss of Alveolar Macrophages and phenotypic changes in COVID-19 lung microenvironment

To characterize immune cell dynamics in the local lung microenvironment, we obtained bronchoalveolar lavage (BAL) samples from 7 Severe COVID-19 with EOD, intubated COVID^+^ patients, 6 COVID^-^ controls, and 5 convalescent patients (i.e. patients who recovered from COVID-19) and performed scRNAseq (**Table S4**). Similar to what we found in circulation, we identified a cluster of S100A12^hi^ Monocytes and a cluster of inflammatory IL-1β^+^ Monocytes that also expressed high levels of inflammatory cytokines *IL1B, CCL3* and *CCL4* (**Fig. 4A-B**). Early phase MoMΦ, expressed higher levels of MoMΦ associated genes *SGK1, MAFB, TREM2*, and *GPNMB* relative to late phase MoMΦ, and were found at significantly increased levels in COVID^+^ patients compared to COVID^-^ and convalescent patients. Late phase MoMΦ expressed higher levels of AM associated genes (e.g. *MARCO, FABP4*), indicating further differentiation toward a resident tissue macrophage (RTM) phenotype. Importantly, we found that AM, the RTM of the lung, were significantly decreased in COVID^+^ patients compared to COVID^-^ patients, but restored to homeostatic levels in convalescent patients. When stratified by age, single cell analysis of normal lung tissue from a cohort of untreated early stage non-small cell lung cancer patients showed a significant decrease of AM in older (>70 years old) patients and an increase in inflammatory MoMΦ, therefore indicating baseline differences in lung MNP composition in elderly populations (**Fig. S6A**)*(45)*.

**Figure 4:**
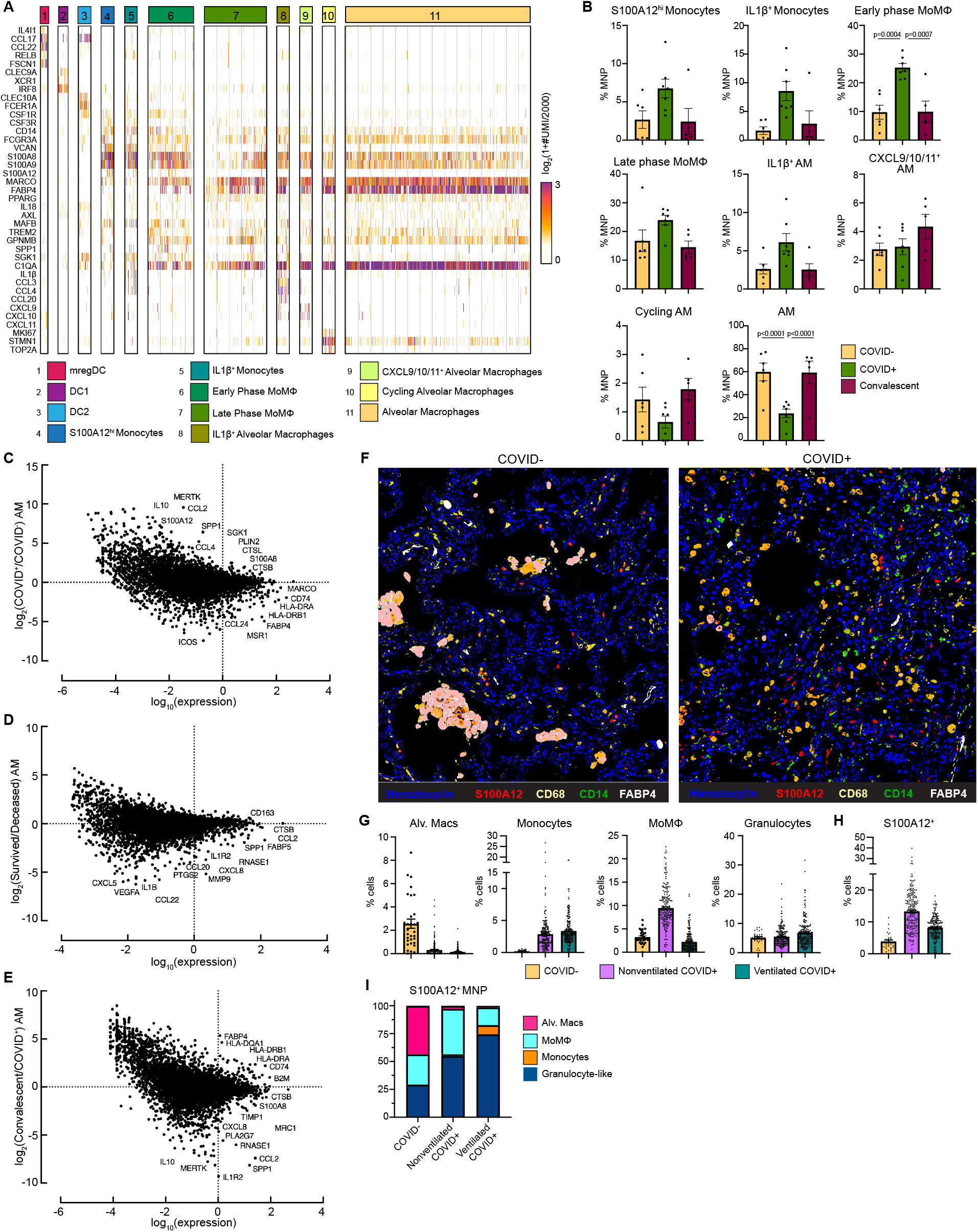
Alveolar Macrophage loss and phenotypic changes in the COVID-19 lung microenvironment. **(A)** Heatmap showing UMI counts of selected genes from myeloid cell scRNAseq clusters from BAL. **(B)** scRNAseq cluster cell frequencies as % mononuclear phagocytes (MNP) in COVID^-^, COVID^+^, or convalescent patient BAL. Differential gene expression between **(C)** AM from COVID^+^ and COVID^-^ patients, **(D)** AM from patients that survived vs deceased, **(E)** AM from convalescent and COVID^+^ patients. **(F)** Overlaid, pseudocolored MICSSS image of COVID^+^ and COVID^-^ lungs, staining for S100A12, CD68, CD14, FABP4, and Hematoxylin. **(G)** Quantification of myeloid cells in MICSSS images, shown as % of cells. AM defined as FABp4^+^CD68^+^ cells; Monocytes defined as CD14^+^ cells; MoMΦ defined as CD14+CD68+ cells; Granulocyte-like cells defined as CD66b^+^ cells or by hematoxylin staining and morphology. **(H)** Quantification of S100A12^+^ cells in MICSSS images, shown as % cells in COVID^-^ patient (n=1), nonventilated COVID^+^ patients (n=2), or ventilated COVID^+^ patients (n=2). **(I)** Distribution of S100A12^+^ cells by cell type in COVID^-^, nonventilated, or ventilated COVID^+^ lungs. For bar graphs, each dot represents patient sample (**B**) or quantification of single MICSSS region of interest (ROI) (**G-H**). Statistical significance (**B**) determined by 2-way ANOVA with Holm-Sidak multiple comparisons correction. Adjusted p-values shown.

In addition to decreased numbers, AM from COVID^+^ patients also expressed higher levels of receptor for advanced glycation end products (RAGE) ligands [*S100A12* (EN-RAGE), *S100A8*] and monocyte chemokines *CCL2* and *CCL4*, while AM from COVID^-^ patients expressed higher levels of Ag presentation genes (e.g. *HLA-DRA, HLA-DRB1, CD74*) and canonical AM markers (*MARCO, MSR1*, and *FABP4*) (**Fig. 4C**). These data suggested that AM from COVID^+^ patients contributed to increased inflammation, recruited monocytes from circulation, and were less proficient at Ag presentation. Further comparison showed that AM from deceased COVID^+^ patients had higher expression of neutrophil chemokines (e.g. *CXCL5, CXCL8*), monocyte chemokine *CCL2*, and inflammatory cytokines (e.g. *IL1B, CCL22*) compared to AM from COVID^+^ patients who survived (**Fig. 4D**). Previous reports have also shown a decline in AM phagocytosis, Ag presentation, and wound healing ability in macrophages with increased *age(46, 47)*. Comparison of AM from convalescent and COVID^+^ patients showed higher expression of class I and II Ag presentation genes after recovery from COVID, thereby suggesting that AM numbers and functionality were restored to baseline in patients who recover from SARS-CoV-2 infection (**Fig. 4E**). In support of this, we did not find any gene expression differences between COVID^-^ and convalescent AM (data not shown).

Spearman correlation analysis of BAL scRNAseq populations showed that the number of inflammatory myeloid cells, the S100A12^hi^ monocytes, IL-1β^+^ Monocytes, MoMΦ, and IL-1β^+^ AM were positively correlated with each other, and negatively correlated with the numbers of AM, Cytotoxic T cells, and Tregs found in BAL (**Fig. S6B-C**). We also observed trending decreases in Tregs in COVID^+^ patients compared to COVID^-^ and convalescent patients, suggesting that hyperinflammation in COVID-19 may be in part due to loss of immunosuppressive activity by Tregs (**Fig. S6D**). All together, these data suggest that the loss of AM, either due to excess inflammation or direct SARS-CoV-2 infection, and their decreased ability to present Ag and recruit and prime T cells may contribute to uncontrolled viral replication and tissue damage*(48)*. Furthermore, elderly patients may be predisposed to more severe disease due to both decreased AM numbers and functionality, and increased inflammatory MoMΦ in the lung.

We confirmed these findings on autopsy lung samples from COVID^+^ patients obtained 10.1±6.2 hours post-mortem using multiplexed immunohistochemical consecutive staining on single slide (MICSSS) (**Table S5**)*(49)*. We observed significant depletion of AM and significant accumulation of CD14^+^ monocytes, CD14^+^ CD68^+^ MoMΦ, and CD66b^+^ granulocyte-like cell infiltration in COVID^+^ lungs compared to a COVID^-^ lung autopsy control obtained from an organ donor (**Fig. 4F-G, S6E**). Comparison between COVID^-^ and COVID^+^ patients also showed increased frequencies of EN-RAGE/S100A12^+^ cells. These changes were not due to ventilation, as similar results were found in ventilated and non-ventilated patients. We also observed a shift in the production of S100A12 from AM in the alveolar air spaces of COVID^-^ lungs, to monocytes, MoMΦ, and granulocyte-like cells in the lung interstitium (**Fig. 4I**). In line with our scRNAseq findings, COVID^+^ lungs from both nonventilated and ventilated patients had decreased Tregs compared to control (**Fig. S6F**). Thus, the loss of AM and Tregs in COVID-19 lungs may lead to an inability to resolve inflammation and initiate tissue repair even after virus is cleared, leading to autonomous inflammation that contributes to morbidity.

## DISCUSSION

In this work, we present systemic and lung high dimensional immunophenotyping on one of the largest single center COVID-19 cohorts to date which was collected during the height of the COVID-19 pandemic in New York City. Here we show that patients with moderate disease had increased numbers of circulating DC, effector and cytotoxic T cells, increased levels of cytokines associated with APC function, and reduced levels of cytokines associated with monocyte and neutrophil activation, mucosal damage, and hyperinflammation. On the other end of the spectrum, severely ill patients had reduced DC, effector and cytotoxic T cells, lower levels of cytokines associated with APC function, and were enriched in immature inflammatory monocytes producing S100A12. Severely ill patients also had high levels of cytokines associated with monocyte and neutrophil activation, mucosal damage, and hyperinflammation.

Notably, we found that lung tissue resident AM were profoundly altered in numbers and functionality in severe COVID-19. AM from COVID^+^ patients expressed higher levels of inflammatory cytokines and decreased levels of HLA class I/II genes compared to AM from COVID^-^ patients, indicating not only a decrease in AM numbers, but also a potential change in their Ag presentation function. AM from deceased COVID^+^ patients were also more inflammatory and expressed higher levels of neutrophil and monocyte attracting chemokines compared to AM from surviving patients, thereby implicating a role for AM in tissue damaging inflammation and recruitment of immature inflammatory myeloid cells from the periphery. All together, these data may suggest that a defect in antigen presentation by altered AM and reduced DC along with mobilization and recruitment of inflammatory monocytes as drivers of disease severity.

The depletion and alteration of the AM pool may be a consequence of direct infection by severe acute respiratory syndrome coronavirus 2 (SARS-CoV-2) and subsequent activation of inflammatory pathways *(48)*. However, alveolar type II cells, the primary angiotensin converting enzyme 2 expressing cells in the lung and target of SARS-CoV-2, also produce granulocyte-macrophage colony stimulating factor (GM-CSF) and have a nonredundant role in maintaining AM in the lung*(50)*. Thus, AM depletion may be multifactorial due not only to inflammation or SARS-CoV-2 infection, but also from decreased GM-CSF in the alveolar milieu. In addition to their role as the first responders to pathogens in the lung, AM play a key role in lung homeostasis, resolution of inflammation, and tissue repair*(51)*. This may explain why elderly patients, who at baseline have decreased AM and increased numbers of inflammatory MoMΦ, are predisposed to increased disease severity.

During development and in homeostatic conditions, RAGE signaling in type I alveolar cells helps maintain alveolar architecture and lung compliance, but is also a known activator of NF-κB signaling*(52)*. Loss of AM derived RAGE ligands such as S100A8/A9 and S100A12 in alveoli may contribute to an inability to maintain proper gas exchange within alveoli. Furthermore, the shift of RAGE ligand production to infiltrating monocytes and Mo-Macs in the lung interstitium we observed in autopsy lungs may exacerbate lung injury, vascular leakage, and lead to increased immature myeloid cell recruitment and infiltration. Increased inflammation has also been implicated in defective transdifferentiation of AT2 cells to AT1 cells, leading to an inability to re-epithelialize and maintain alveolar barrier integrity*(53)*. Our data suggest that maintaining and restoring AM numbers early during infection may be a valid therapeutic strategy that may protect airway integrity and initiate an early innate and adaptive immune response, while limiting the expansion and recruitment of immature inflammatory myeloid cells from the periphery*(54)*.

## MATERIALS AND METHODS

### Study Design

The goal of our study was to identify drivers of COVID-19 severity and death in order to support the identification and development of tailored immunotherapy strategies to halt disease progression. To do so, we performed high dimensional immunophenotyping on hospitalized COVID-19 patients at the Mount Sinai Hospital from March 2020-December 2020. We characterized inflammation patterns using the Olink platform which allowed us to detect 92 different proteins from patient sera. To understand the diversity of immune patterns, we performed unbiased clustering analysis and noted immune patterns that correlated with disease severity, comorbidities, and patient outcome. We grouped immune patterns based on these clinical parameters and calculated protein module scores based on the covariance patterns of different cytokines. Next, we characterized circulating immune cells using CyTOF on whole blood samples and scRNAseq on PBMC. Here, we used unbiased clustering on the PBMC scRNAseq to identify immune cell populations and compared the frequencies across Olink groups. Following this, we integrated our seromics data with scRNAseq to identify 4 distinct immune responses to COVID-19. To characterize local changes to the lung immune microenvironment, we obtained BAL samples from COVID^+^, COVID^-^, and convalescent patients and performed scRNAseq. We further expanded our characterization of the lung using MICSSS on lung autopsy samples and quantified changes in myeloid cell infiltration.

### Mount Sinai COVID-19 Biobank

Electronic medical records (EMR) from patients admitted to the Mount Sinai Hospital for suspected or confirmed COVID-19 were screened each morning by a team of volunteers and team physicians for enrollment into the Mount Sinai COVID-19 Biobank. Due to the difficulty of obtaining direct informed consent during the pandemic, the Institutional Review Board (IRB) approved sample collection from patients before consent was obtained. Patients were made aware of planned sample collection with documents provided during hospital registration and provided instructions for opting out. Patient consent was subsequently obtained by contacting patients via hospital-room phone, phone call after discharge, or through legally authorized representatives after death of patients. All patient samples used and presented in these analyses were from consented patients. This study was approved by the Institutional Review Board of the Mount Sinai School of Medicine under IRB-20-03276. EMR and deidentified clinical data for each patient was pulled from Epic electronic health record using Epic Hyperspace, Epic Clarity, and the Mount Sinai Data Warehouse, and summarized per 24 hour period measures throughout length of hospital stay. Period window was defined as 12 PM to 12 PM to match blood draw time.

### Clinical Blood Sample Collection and Processing

At each collection timepoint, patient serum was collected in a Vacutainer^®^ Plus Plastic SST^™^ Blood Collection Tubes with Polymer Gel for Serum Separation tube. 2x BD Vacutainer^®^ CPT^™^ Cell Preparation Tube with Sodium Heparin were collected for Plasma and PBMC collection. Samples were obtained by nurses or phlebotomists as part of clinical care, collected from hospital floors by “Running Team” volunteers, and delivered to the laboratory for processing. Blood samples were kept on gentle agitation at room temperature (RT) and processed by Blood Processing Team volunteers of the Mount Sinai COVID-19 Biobank in Biosafety level 2 plus (BSL-2+) facilities on the day of collection.

SST tubes were centrifuged at RT at 1300 relative centrifugal force (rcf) for 10 minutes (mins) and serum was banked into cryovials for storage in liquid nitrogen (LN). Whole blood samples for CyTOF were taken from CPT tubes and directly stained with a lyophilized antibody panel using Fluidigm MaxPar Direct Immune Profiling Assay (MDIPA) tubes for 30 mins at RT. Stained whole blood samples were then stabilized and fixed with Prot1 Proteomic Stabilizer for 10 mins at RT before storage at −80°C as previously described*(55)*. CPT tubes were then centrifuged at 1800 rcf for 15 mins to separate plasma and PBMC. Plasma was aspirated and banked into cryo-vials for storage in LN. The PBMC cell layer was collected, washed with phosphate buffered saline (PBS) and collected by centrifugation at 300 rcf for 15 mins. Cell viability and counts were assessed by acridine orange and propidium iodide (AOPI) staining in automated Nexcelom Cellometer Cell Counters. PBMC were resuspended at a concentration of ~10×10^6^ cells/mL in Human Serum Ab and 10% dimethyl sulfoxide (DMSO) and stored at −80°C for 24 hours before transfer to LN storage.

### Olink measurements of COVID-19 serum, data normalization, and clustering analysis

Olink was performed on COVID-19 patient serum samples in BSL2^+^ according to manufacturer instructions. Count (Ct) values were generated by Olink NPX manager software. To control for technical variability between plates, we included in each plate 2 technical control replicates from a single mixture of pooled blood from healthy donors and estimated a control value per plate, defined as:

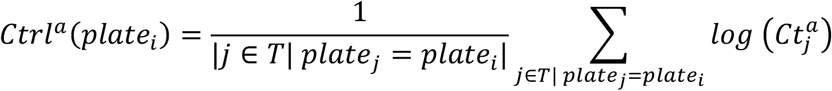

Where: a is a given analyte, i is a given sample, T is the set of technical control replicates, plate_i_ is the plate of sample i, and 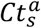 is the raw Olink Ct value of analyte a in sample s.

Normalized Ct values 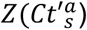 were definied as the z-scores of the plate-adjusted, log transformed Ct values 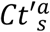:

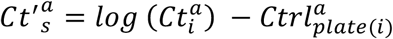

Samples with similar normalized Ct profiles 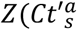 were clustered using Kmeans++ (https://github.com/tanaylab/tglkmeans) with k=15.

Olink protein module scores were calculated by averaging the normalized z-scores of each module analyte.

### CyTOF Data acquisition and analysis

MDIPA stained whole blood samples were thawed using the SmartTube Prot 1 Thaw/Erythrocyte Lysis protocol. Samples were subsequently barcoded and pooled utilizing the Fluidigm Cell-ID 20-Plex Pd Barcoding Kit and stained with an antibody cocktail against fixation stable markers for more in depth immune profiling. Following sample barcoding and staining, samples were fixed with 2.4% paraformaldehyde in PBS with 0.08% saponin and 125 nM Iridium (Ir) for 30 mins at RT and stored in Cell Staining Buffer until acquisition. Immediately prior to data acquisition, samples were washed once in Cell Acquisition Solution and resuspended at a concentration of 1*10^6 cells/mL for acquisition (including 10% Fluidigm EQ Normalization Beads). The resuspended cells were then acquired on the Helios Mass Cytometer supplemented with a wide bore injector at an event rate <400 events/second. After data acquisition, samples were debarcoded using the Astrolabe Diagnostics platform. Cell populations were identified by a combination of an automated approach using the Astrolabe Diagnostics Platform and manual gating as previously described*(55)*.

### PBMC preparation for scRNAseq

PBMC samples were selected based on manual EMR chart review taking into account, and controlling for patient demographics and treatments. Frozen PBMC were thawed at 37°C and resuspended in RPMI media+ 10% fetal bovine serum (FBS) with 25 U/mL Benzonase before centrifugation at 350 rcf for 5 mins. Cells were resuspended in media and viable cells were counted by AOPI staining in Nexcelom Cellometer Cell Counters. Combinatorial hashes were prepared in wash buffer (PBS + 0.5% bovine serum albumin (BSA)). 500,000 live cells were stained with hashes for 20 mins on ice before 3 washes in wash buffer. Cells were filtered through a 70 μm filter and then a 40 μm filter twice. Filtered cells were counted and loaded with a targeted cell recover of 35,000 cells/lane across 8 lanes of 5’ v1.1 NextGEM assay.

### Patient selection for BAL

Respiratory samples for research were allocated from BAL obtained from patients ≧ 18 years of age undergoing bronchoscopy with BAL fluid collection for clinical reasons. Patient groups included the following: (1) positive SARS-CoV-2 PCR with COVID-19 related acute respiratory failure requiring intubation and mechanical ventilation; and (2) negative SARS-CoV-2 PCR with suspected lung cancer. Patients in Group 2 who had previous positive SARS-CoV-2 PCR or SARS-CoV-2 Ab were designated as COVID-19 convalescent. Informed consent for bronchoscopy with BAL fluid collection was obtained separately from consent for research.

#### Group 1

Patients were identified by the attending critical care physician providing clinical care. Patients were intubated at the discretion of the critical care team for progressive respiratory failure, as evident by worsening hypoxemia, hypercapnia, or work of breathing despite support by high-flow nasal cannula oxygen or non-invasive ventilation. Bronchoscopy with BAL fluid collection was performed within 72 hours of first intubation if clinically indicated. At our center, all patients with COVID-19, including those with COVID-19-related respiratory failure, were managed according to guidelines developed and updated by the Mount Sinai Health System as new data regarding care of patients with COVID-19 became available*(56)*. Patients requiring intubation and mechanical ventilation for COVID-19-related respiratory failure were additionally managed with a low tidal volume ventilation strategy*(57)*.

#### Group 2

Patients were identified by the pulmonologist and physician assistant who performed the bronchoscopy with BAL fluid collection, which was carried out for suspected lung cancer requiring a diagnostic and staging procedure. Negative SARS-CoV-2 PCR was obtained 2-5 days prior to bronchoscopy. Patients were selected on the basis of suspected lung cancer not greater than 5 cm in greatest dimension. Patients with known history of, or clinical suspicion for, active infectious or inflammatory lung diseases were excluded.

### BAL fluid collection

All respiratory specimens were collected using sterile, flexible, fiberoptic bronchoscopes. Bronchoscopes were flushed prior to the procedure with 5 mL sterile saline, which was collected for research. For Group 1, a single-use bronchoscope (Ambu) was inserted through the endotracheal tube. For Group 2, a reusable bronchoscope was inserted through a laryngeal mask airway or endotracheal tube placed for the procedure. After airway inspection, the bronchoscope was wedged in a distal airway of interest selected by pre-procedure imaging. Sterile saline was instilled in 30 mL aliquots (up to 90 and 210 mL for Groups 1 and 2, respectively) and aspirated. Aspirated BAL fluid was split into parts for clinical use, which included fluid sent for clinical microbiological analysis, and research use, which was transported immediately to the research laboratory on ice and processed as below.

### BAL fluid processing

BAL samples were processed within 30 mins of sample collection. Collected BAL was filtered twice through 70 μM filters and centrifuged at 350 rcf for 5 mins at 4°C. BAL supernatant was collected and treated with 0.1% Triton X-100 for 1 hour to inactivate virus before aliquoting into cryovials for storage at −80°C. BAL cells were incubated with Red Blood Cell Lysis buffer (Fisher Scientific) for 5 mins at RT before washing with PBS+ 0.5% BSA and centrifugation at 350 rcf for 5 mins at 4°C. Viable cells were counted by AOPI staining in Nexcelom Cellometer Cell Counters. 2 lanes of 8000 cells from each sample were loaded onto the 10x Chromium Controller for scRNAseq, 5’ v1.1 NextGEM assay.

### PBMC and BAL scRNAseq data processing

For all scRNAseq datasets, debris/empty droplets were identified with cells that had gene expression (GEX) UMI counts<22. Cells were identified by finding local minimum in the GEX UMI distribution, keeping only cells ≧ the local minimum UMI count. On average, this was 371 UMI counts. For PBMC scRNAseq, hashtag oligo (HTO) UMI counts <22 and excluded from further analysis. For HTO transformation, each feature in the HTO matrix (linear space) was subtracted from its 5^th^ quantile and divided by its 95^th^ quantile. Each cell was subsequently divided by its UMI sum. Because hashing reads were not consistently detected, we underwent additional processing to dehash cells.

#### Initial mapping of cells to samples using only HTOs

A “key” matrix with biological samples as rows and HTO features as columns was created. Each unit in the matrix was populated with a value of “1” if the sample was supposed to be positive for the hashtag and “0” if it was not. For each cell, pairwise distances with a cosine similarity metric were computed from the key matrix to generate a cosine similarity matrix. Samples with the highest cosine similarity could then be assigned to the cell. Initially assigned mappings were called “hto-ini”.

Additionally, a signal to noise ratio (SN) was for each cell was calculated by subtracting the highest cosine similarity from the second highest cosine similarity. The SN ratios for each initially assigned sample usually followed a bimodal distribution. The relative minima of the parametric density of SN ratios was used in order identify the local minimum of this distribution. Only cells with SN ≧ the local minimum were kept as the final cells belonging to that sample. We call these final mappings “sample-hto”

#### Initial mapping of cells to samples using only HTOs and Souporcell

We used Souporcell to cluster the cells based on the polymorphisms detected from the RNA-Seq alignment*(58)*. We also inputted the genotypes inferred from whole genome DNA-Seq data as a reference in Souporcell in order to map the clusters to respective patients. Next, we leveraged the Souporcell subject mappings along with the initially assigned HTO mappings (hto-ini) in order to deconvolute the patient to the various timepoints. For patients with single timepoints, we assigned the entire Souporcell cluster to the sample. For patients with multiple timepoints, we performed the hashtag de-multiplexing strategy described above to map the cells from the cluster to the patient’s timepoints. We called these mapped cells “sample-soc-hto”.

#### Final mappings

We intersect the output from both strategies above (“sample-hto” and “sample-soc-hto”) in order to get a consensus cell-sample mapping. Only these consensus cells were used for further downstream analysis.

### scRNAseq Analysis

Briefly, mRNA reads were tagged with a cell barcode and UMI. These reads were aligned, and count matrices were built. Cell barcodes with at least 500 UMIs were extracted, and cells that were comprised of more than 25% of reads from the mitochondrial genome were filtered from subsequent analysis. The variability in cell counts or UMI counts across samples were not confounding variables in downstream analyses. The R package Seurat was then used for data scaling, clustering, dimensionality reduction, and downstream differential gene expression analyses*(59–62)*. The function SCTransform was used to scale and identify variable genes that constituted principal components for principal component analysis (PCA). The first 15 principal components were used to perform UMAP reduction, once a shared nearest-neighbor graph had been generated and clustering had been performed based on the Louvain method for community detection. Cells were down-sampled to 2000 UMIs per cell and variable genes were selected. Gene module analysis was performed by computing a Pearson correlation matrix between genes for each sample using the R package scDissector*(63)*. Highly correlated genes were grouped into gene modules by hierarchical clustering. These modules were used to determine cellular identity for each cluster that was present across samples.

scRNAseq immune cell cluster frequency correlations, and integrated scRNAseq cell frequencies and Olink proteomics were calculated using the corrplot package (v0.88) and visualized using the pheatmap package (v1.0.12) in R.

### Lung Autopsy Tissue Section Preparation

Lung autopsy samples were collected within 24 hours of death (average 10.1±6.2 hours) and fixed in 10% neutral-buffered formalin for 24 hours before transfer to 70% Ethanol (EtOH). Samples were then embedded in paraffin and 4 μM tissue sections formalin fixed paraffin embedded (FFPE) sections were cut onto glass slides and baked at 37°C overnight.

#### Autostainer (Bond Rx, Leica Biosystems)

Slides were covered with covertiles (Bond Universal Covertiles, Leica biosystems) and baked for 10 mins at 57°C. Slides were deparaffinizied in dewax solution and rehydrated in decreasing concentrations of EtOH. Tissue sections were then incubated in Ag retrieval solution (pH 6 or 9) at 95°C for 20 mins. Tissue sections were incubated in 3% hydrogen peroxide (Bond Polymer Refine Detection Kit DS9800, Leica Biosystems) for 15 mins to block endogenous peroxidases. Next, tissue sections were incubated in serum-free protein block solution (Dako X0909) for 30 mins to block nonspecific antibody binding. After the first staining cycle, Fab fragments (AffiniPure Fab Fragment Donkey anti-mouse (715-007-003) or anti-rabbit IgG (711-007-003)) against that primary antibody species were used to block carryover staining whenever there was a repeat of same primary antibody species. Primary antibody staining was performed for 1 hour at room temperature followed by secondary antibody staining. Polymer detection system (Bond Polymer Refine Detection Kit #DS9800, Leica Biosystems) was used for horseradish peroxidase signal amplification. Chromogenic revelation was performed using ImmPact AEC (3-amiino99-ethylcarbazole) substrate (Vector Laboratories, SK4205) for preset incubation times. Slides were counterstained with hematoxylin (Bond Polymer Refine Detection Kit, DS9800, Leica Biosystems.

#### Manual staining

Slides were mounted with a glycerol-based mounting medium (Dako, C0563) and scanned for digital imaging (Hamamatsu NanoZoomer S60 Whole Slide Scanner). Same slides were successively stained, as per MICSSS protocol*(49)*. Coverslips were removed by placing slides in a rack and immersing in hot tap water at 56°C until mounting media dissolved. Chemical destaining between stains was performed by immersing slides in gradually diluted EtOH solutions.

### MICSSS coexpression analysis

To analyze marker coexpression, a pseudofluorescence composite image of all chromogenic markers was created. The same region of interest (ROI) were selected from images of each marker in QuPath (https://qupath.github.io/, Bankhead et al. 2017) and exported as PNG formatted images without downsampling. Images of different immunostains belonging to the same ROI were transferred to Fiji-ImageJ and co-registered using the TrakEM2 plug-in*(64, 65)*. Color deconvolution was performed using H-AEC vectors for each image to split the RGB images into three 8-bit channels including hematoxylin (blue), AEC (red chromogen color), and residual (green) channel. The best hematoxylin channel was selected as the nuclear channel. AEC channels representing staining of each marker were assigned to different colors by using the lookup tables (LUTs) function of Fiji while hematoxylin channel was assigned to blue color to mimic fluorescent DAPI staining. Next, color inversion was done on all channels and then merged to achieve a multiplexed pseudofluorescent image. We optimized brightness and contrast settings to facilitate visualization for each immunostain channel by comparison with original chromogen images but did not change underlying image pixel values for quantification.

### MICSSS quantification

Stained images were scanned at 40x resolution into the .ndpi format and uploaded to Amazon Web Services super computer clusters for high-speed analysis on a Python-based Anaconda Jupyter Notebook. Raw red-green-blue (RGB) thumbnail 1.25x resolution images were analyzed using an in-house tissue recognition algorithm that enhanced tissue contours and used optical densities of pooled pixels across the image to determine a tissue mask. The image was then rigidly registered with the corresponding images across all markers for the same tissue within the MICSSS panel. This registration used a third party SimpleElastic package for Python (https://simpleelastix.readthedocs.io/RigidRegistration.html). Following linear registration, the highest resolution image for each marker (40x) was spliced into multiple tiles that spanned approximately 2000 μm in each dimension with 20% surface area overlap in each direction. Each tile was then denoted with an X,Y pooled pixel coordinate value so that the appropriate corresponding tiles across all the markers in the panel for the same tissue would be analyzed together. Each set of tiles was deconvoluted to extract the hemoxylin channel, which remained consistent across all markers. The hematoxylin channel was then registered with an affine registration (which accounts for shear, scale, rotation, and translational dislocation) and a “b-spline” elastic warping to account for any local tissue warping or tissue damage (https://simpleelastix.readthedocs.io/NonRigidRegistration.html)

The vector field transformation matrix produced from the high-resolution affine and b-spline registrations was then applied to the raw RGB tile. Registered RGB tiles were analyzed in parallel across the multiple cores of the AWS supercomputer, trimmed to eliminate overlap, and concatenated to produce one final elastically registered RGB image per marker. 100 ROIs of about 500×500 μm were randomly chosen in the image based on where tissue resided. These were chosen from the last tissue mask in the panel to account for any tissue damage or warping. Each of these ROI was processed in parallel across the multiple cores in the AWS supercomputer. Next, we used the Stardist package for Python (https://github.com/stardist/stardist) that was trained with hematoxylin and eosin staining, to segment each ROI, and to determine the centroids and morphological properties of each determined cell. We previously optimized the sensitivity of this algorithm with these tissues and therefore used an overall sensitivity value of 0.1 for the algorithm. This algorithm provides information for the nucleus of each cell in the ROI, which we then artificially expanded by 5 microns to simulate the cell’s cytoplasm. Overlapping cytoplasm from adjacent cells in dense regions were condensed to prevent overcounting surface area.

Each cell per ROI was then analyzed for marker positivity. ROI were deconvoluted for its AEC detection channel and each cell was translated to a median AEC value from pixels that resided in its nucleus and artificially expanded cytoplasm. If the median AEC value for the cell was above the threshold value for positivity deemed for that marker, the cell was considered positive. Threshold values per marker were determined with a pathologist. % positive cells was determined by the number of cells above the threshold for that marker over the total number of cells in that ROI. Co-expression analysis was performed by determining if a cell was positive for the multiple markers of interest over the total number of cells in that ROI. We plotted each value per ROI.

### Statistical analysis

Data analysis was performed using Prism version 9.2.0 (GraphPad software) or in R version 3.6.3 and presented as stated in figure legends. 2-way ANOVA with Tukey’s multiple comparisons correction was used for Olink comparisons while 2-way ANOVA with Holm-Sidak multiple comparisons correction was used to calculate cell frequency changes for CyTOF and scRNAseq. For spearman correlation coefficients, p<0.05 was considered statistically significant p-values in correlation matrices indicated as *p<0.05, **p<0.005, ***p<0.0005.

## Supplementary Materials

Fig. S1. Unsupervised K-means++ clustering of COVID-19 serum proteomics identified 15 distinct immune patterns

Fig. S2. Clinical characteristics of Olink serum clustering analysis

Fig. S3. Covariance matrix of Olink analytes in COVID-19 serum

Fig. S4. Olink protein module scores are stable and associated with clinical outcome.

Fig. S5. High dimensional characterization of circulating lymphocytes in COVID-19

Fig. S6. High dimensional characterization of immune cells in BAL

Table S1. Mount Sinai COVID-19 Biobank patient summary

Table S2. Mount Sinai COVID-19 disease severity classification

Table S3. Clinical information for PBMC scRNAseq samples.

Table S4. Clinical information for BAL samples.

Table S5. Clinical information for lung autopsy samples.

## Acknowledgements

We would like to thank the patients and their families for agreeing to be a part of this study. We thank all the volunteers of the Mount Sinai COVID-19 Biobank which was comprised of a redeployed workforce by the following centers, programs, departments, and institutes within the Icahn School of Medicine at Mount Sinai: the Human Immune Monitoring Center; Mount Sinai COVID-19 Informatics Center; Program for the Protection of Human Subjects; Department of Psychiatry; Department of Genetics and Genomic Sciences; Department of Medicine; Department of Oncological Sciences; The Precision Immunology Institute; Tisch Cancer Institute; Icahn Institute for Data Science and Genomic Technology; Friedman Brain Institute; Charles Bronfman Institute of Personalized Medicine; Hasso Plattner Institute for Digital Health; Mindich Child Health and Development Institute; and Black Family Stem Cell Institute. We would also like to thank all the front-line healthcare workers and staff who made this work possible.

## Author contributions

Conceptualization: STC, MF, S-KS, AHR, BDB, EK, MM

Methodology: STC, MDP, MB, UC, KM, CAP, JLH, EK, MM

Software: STC, MDP, DMDV, MB, BL, DG, DD, RT, NDB, EK

Validation: STC, MDP

Formal Analysis: STC, MDP, MB, EK

Investigation: STC, MDP, DMDV, MB, AT, BL, DG, DD, TD, RM, KN, RT, ZZ, JLB, CC, HJ, NDB, EK

Resources: DMDV, NWS, KM, NB, RT, ZZ, UC, KM, SA, SB, MR, TH, SK-S, AHR, CAP, JLH

Data Curation: MDP, DMDV, MB, BL, DG, DD, RT, SK-S, NDB, EK

Writing- Original Draft: STC, MDP, MB, AT, DD, EK

Writing-Review and Editing: STC, MDP, MM

Visualization: STC, MDP, AT, EK

Supervision: EK, MM

Project Administration: JLB, CC, SK-S, AHR, SG, AWC, MM,

Funding acquisition: AHR, BDB, SG, AWC, MM

## The Mount Sinai COVID-19 Biobank

The following investigators and volunteers participated in the Mount Sinai COVID-19 Biobank effort including electronic medical record screening, patient consenting, collection kit assembly, nurse outreach, specimen transport, sample processing, biobanking, data acquisition, data management, and data processing: Charuta Agashe, Priyal Agrawal, Alara Akyatan, Kasey Alesso-Carra, Kimberly Argueta, Eziwoma Alibo, Kelvin Alvarez, Angelo Amabile, Steven Ascolillo, Craig Batchelor, Rasheed Bailey, Priya Begani, Grace Chung, Paloma Bravo Correra, Larissa Burka, Sharlene Calarossi, Lena Cambron, Gina Carrara, Serena Chang, Esther Cheng, Jonathan Chien, Mashkura Chowdhury, Jonathan Chung, Cansu Cimen Bozkus, Phillip Comella, Dana Cosgrove, Francesca Cossarini, Liam Cotter, Arpit Dave, Bheesham Dayal, A. Cassandra De Jesus, Maxime Dhainaut, Rebecca Dornfeld, Katie Dul, Melody Eaton, Nissan Eber, Cordelia Elaiho, Frank Fabris, Jeremiah Faith, Dominique Falci, Susie Feng, Brian Fennessy, Marie Fernandes, Nancy Francoeur, Sandeep Gangadharan, Joanna Grabowska, Gavin Gyimesi, Maha Hamdani, Diana Handler, Jocelyn Harris, Matthew Hartnett, Sandra Hatem, Manon Herbinet, Elva Herrera, Arielle Hochman, Gabriel E. Hofman, Laila Horta, Etienne Humblin, Jessica S. Johnson, Gurpawan Kang, Neha Karekar, Subha Karim, Geoffrey Kelly, Jessica Kim, Jose Lacunza, Alana Lansky, Dannielle Lebovitch, Grace Lee, Gyu Ho Lee, Jacky Lee, John Leech, Lauren Lepow, Mike Leventhal, Lora E. Liharska, Katherine Lindblad, Alexandra Livanos, Rosalie Machado, Zafar Mahmood, Kelcey Mar, Shrisha Maskey, Paul Matthews, Katherine Meckel, Saurabh Mehandru, Anthony Mendoza, Cynthia Mercedes, Dara Meyer, Ronaldo Miguel de Real, Gurkan Mollaoglu, Sarah Morris, Emily Moya, Nicole Ng, Marjorie Nisenholtz, George Ofori-Amanfo, Kenan Onel, Merouane Ounadjela, Manishkumar Patel, Vishwendra Patel, Cassandra Pruitt, Shivani Rathi, Jamie Redes, Ivan Reyes-Torres, Alcina Rodrigues, Alfonso Rodriguez, Vladimir Roudko, Evelyn Ruiz, Kevin Rurak, Pearl Scalzo, Ieisha Scott, Pedro Silva, Alessandra Soares Schanoski, Hiyab Stefanos, Meghan Straw, Sayahi Suthakaran, Collin Teague, Kevin Tuballes, Bhaskar Upadhyaya, Verena Van Der Heide, Natalie Vaninov, Daniel Wacker, Laura Walker, Hadley Walsh, C. Matthias Wilk, Lillian Wilkins, Jessica Wilson, Karen M. Wilson, Hui Xie, Luyi Xu, Li Xue, Naa-akomaah Yeboah, Nancy Yi, Mahlet Yishak, Sabina Young, Alex Yu, Nicholas Zaki, Nina Zaks, and Renyuan Zha.

## Data and materials availability

All consented data used will be deposited in public repositories upon publication. Code used for MICSSS quantification available upon request.

