## Supplementary Materials for "Shift of lung macrophage composition is associated with COVID-19 disease severity and recovery"

### Supplementary Material

**A**

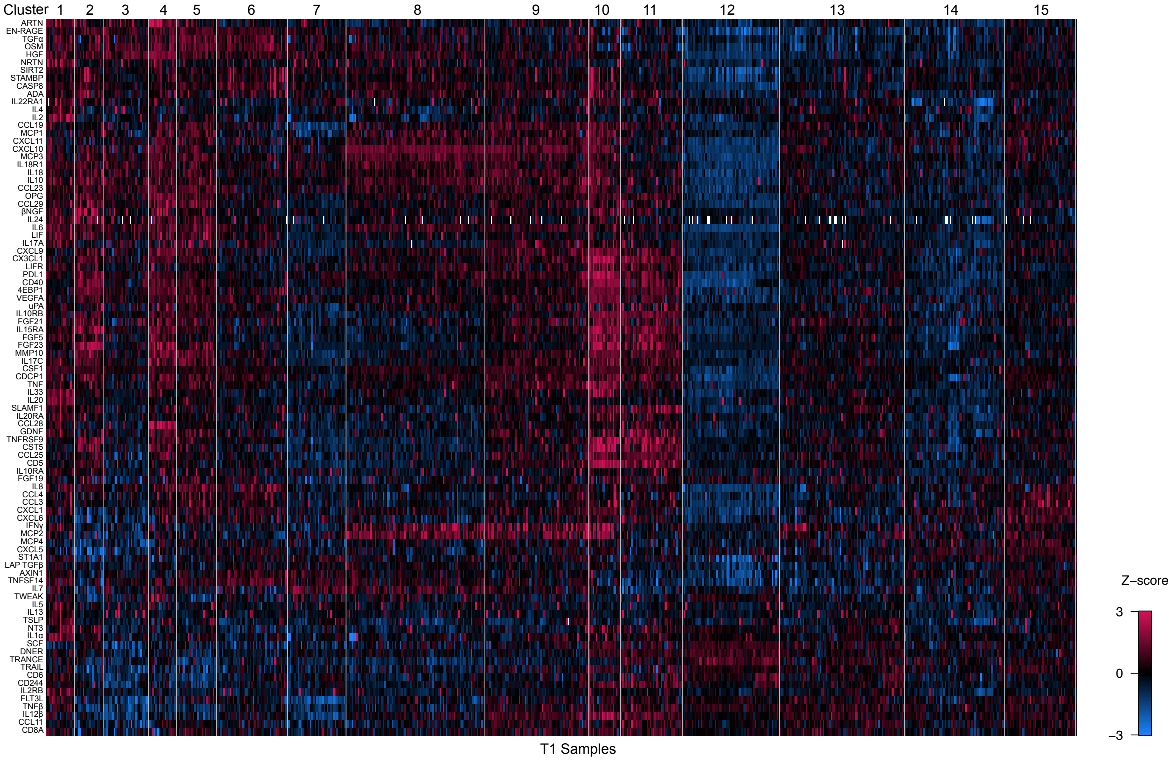

**B**

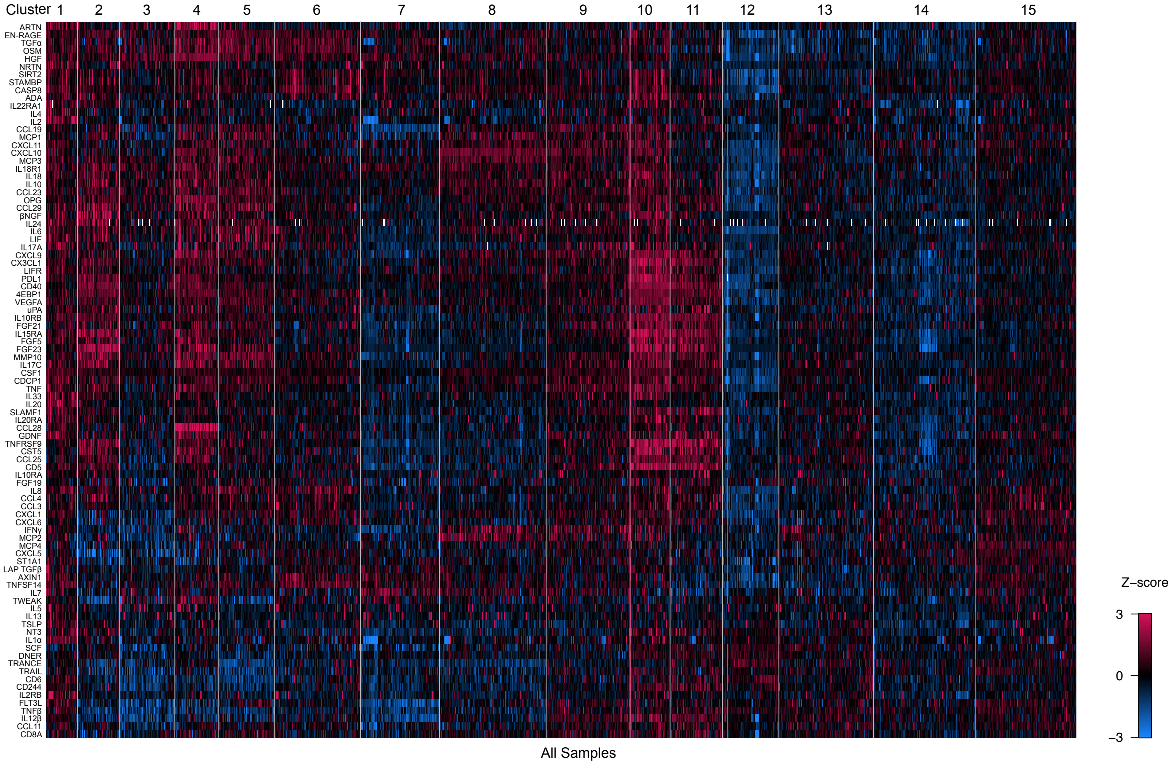

**Supplementary Figure 1. Unsupervised K-means++ clustering of COVID-19 serum proteomics identified 15 distinct immune patterns (A)** Heatmap showing cytokine profiles of COVID-19 sera measured with the Olink inflammation panel. Unsupervised K-means++ clustering was performed on normalized values (z-scores) for all serum samples. 15 clusters were identified. Rows denote each protein measured; columns denote proteomic profile of each patient serum sample. **(B)** Heatmap showing cytokine profiles of COVID-19 sera measured with the Olink inflammation panel. Unsupervised K-means++ clustering was performed on normalized values (z-scores) for all T1 serum samples. 15 clusters were identified. Rows denote each protein measured; columns denote proteomic profile of each patient serum sample.

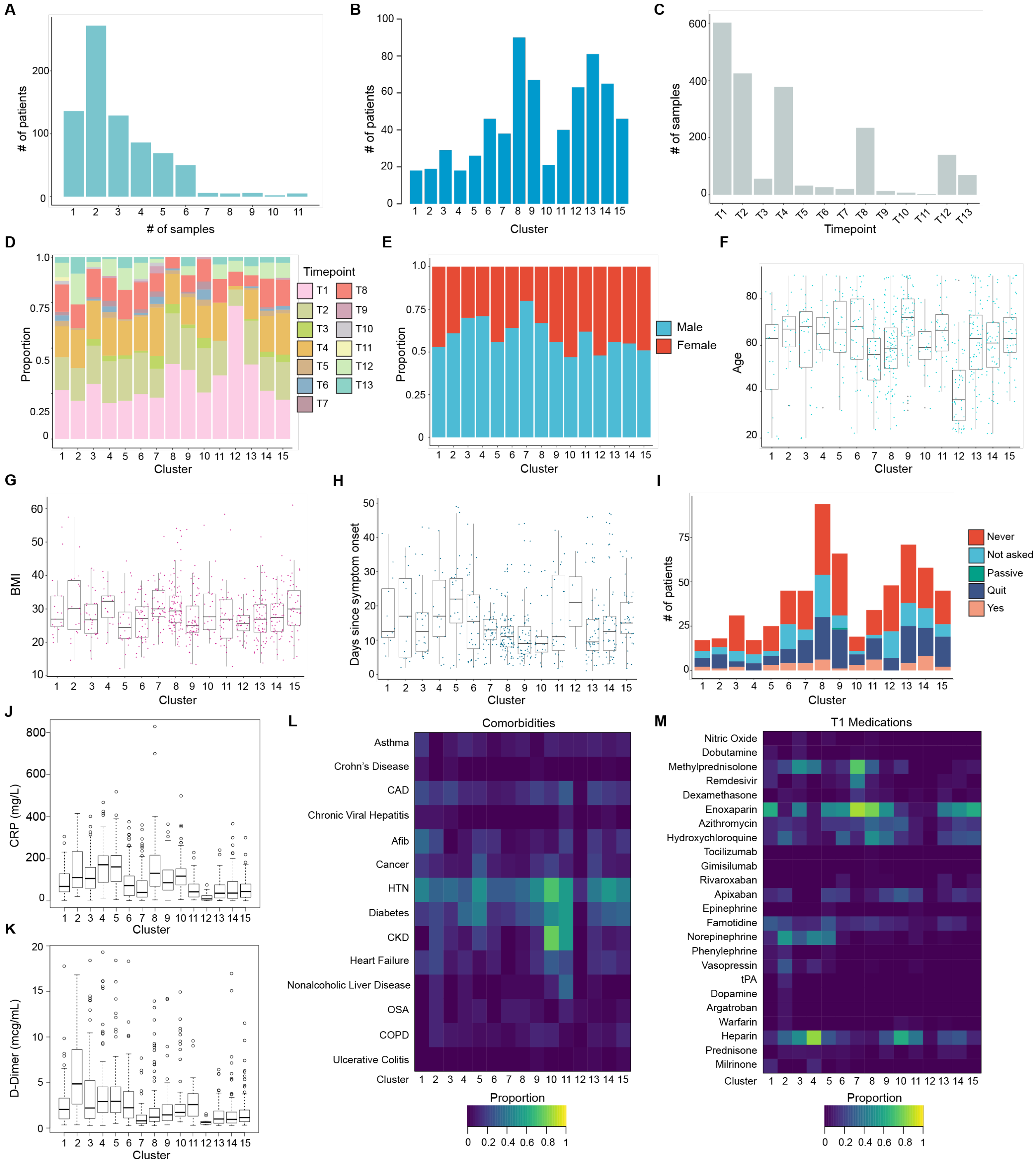

**Supplementary Figure 2. Clinical characteristics of Olink serum clustering analysis. (A)** Histogram showing number of patients by number of samples analyzed. **(B)** Histogram showing number of patients per Olink cluster. **(C)** Histogram showing number of samples acquired for each Timepoint. **(D)** Timepoint distribution per Olink cluster. **(E)** Proportion of sex by Olink cluster. Boxplot showing patients' age **(F)** and BMI **(G)**, and days post symptom onset **(H)** across Olink clusters. Cluster assignment set by first timepoint for each patient. **(I)** Stacked histogram showing smoking status for patients by Olink Cluster. Boxplot showing C-reactive protein **(J)** or D-Dimer levels **(K)** for first available patient samples. Cluster assignment set by first timepoint for each patient. **(L)** Heatmap showing proportion of patients with comorbidities by Olink Cluster. Cluster assignment set by first timepoint for each patient. **(M)** Heatmap showing proportion of patients receiving medications at time of first sampling.

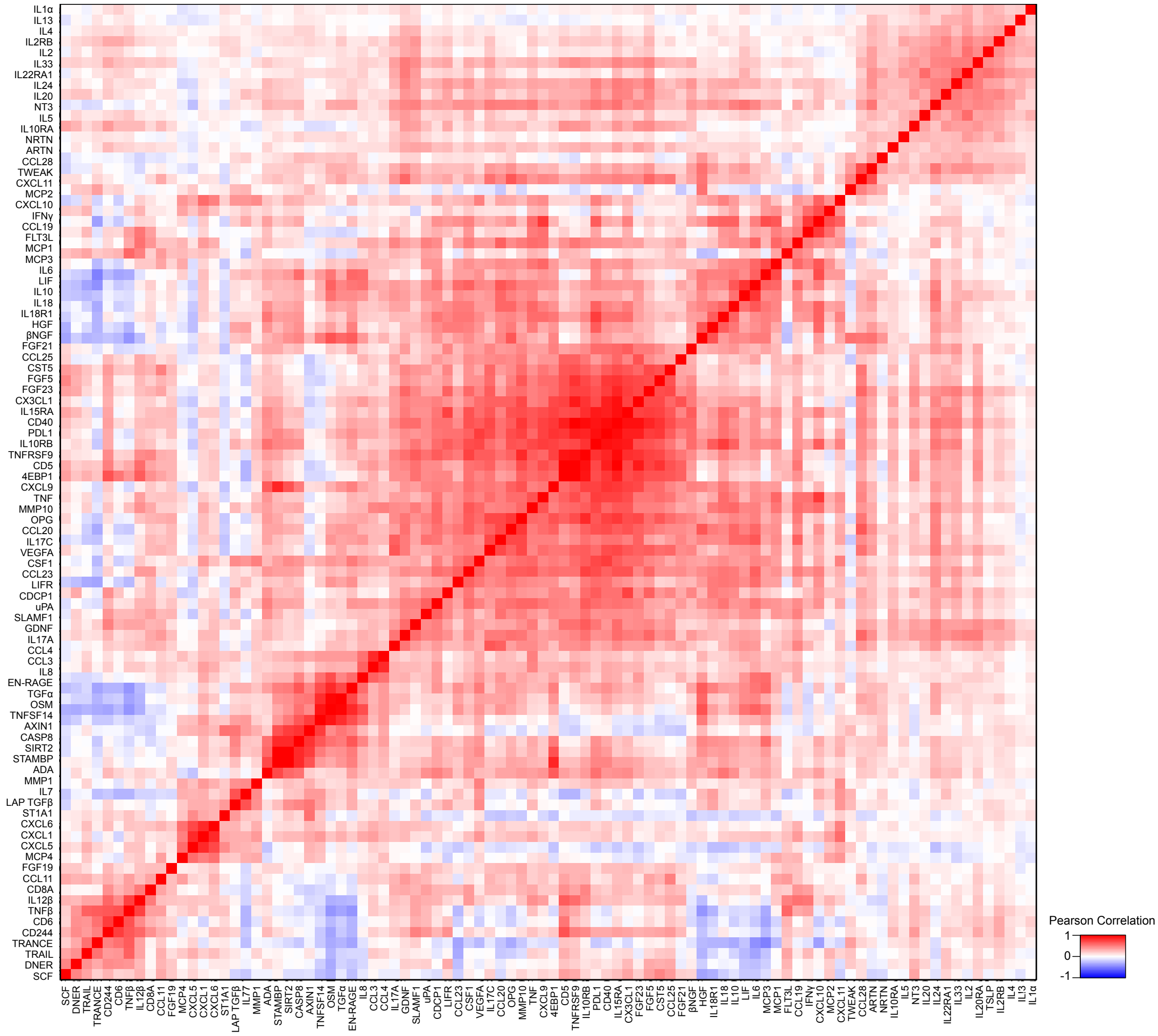

**Supplementary Figure 3. Covariance matrix of Olink analytes in COVID-19 serum.** Heatmap showing Pearson correlations between Olink analytes for all COVID-19 patient serum samples.

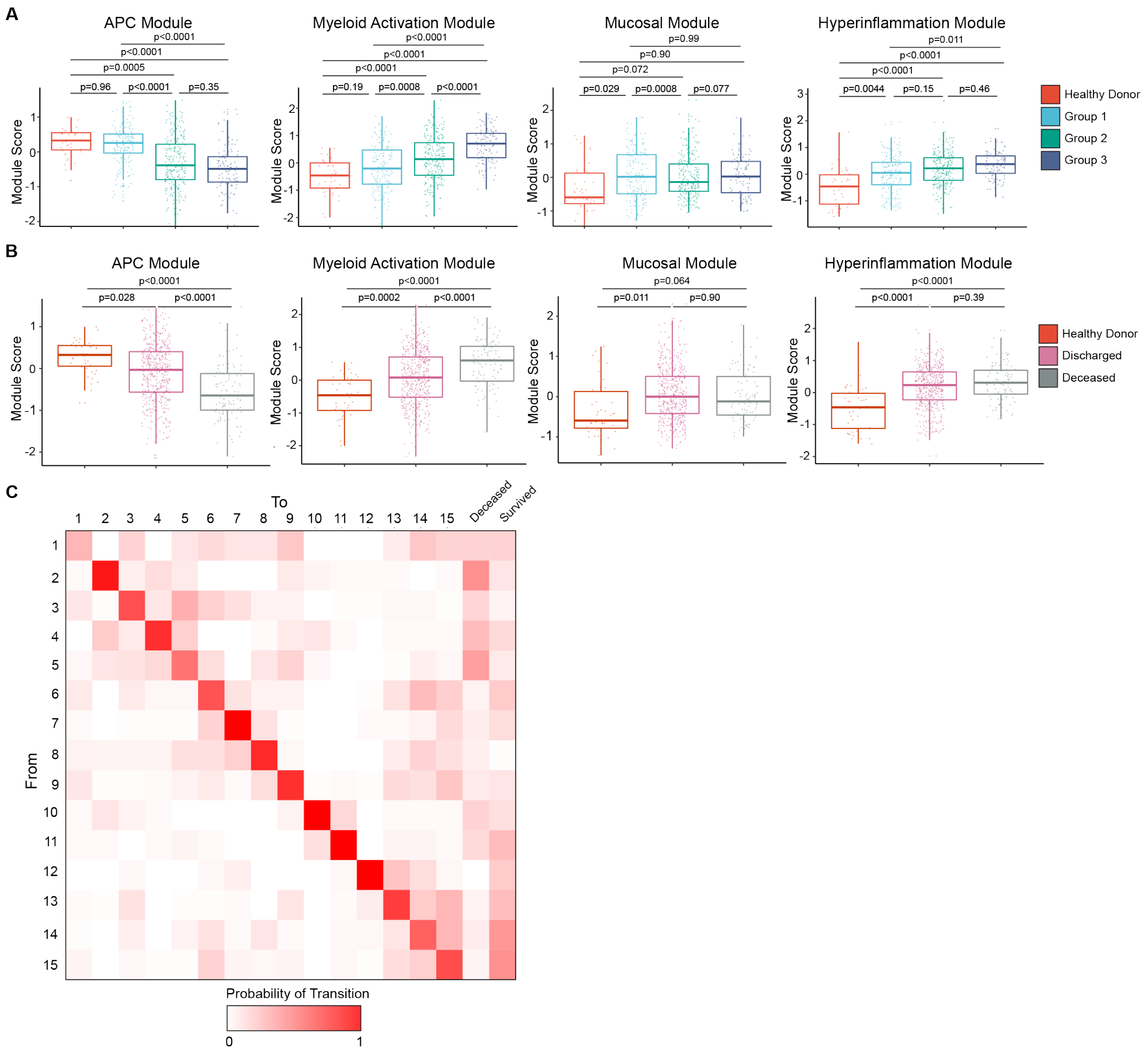

**Supplementary Figure 4. Olink protein module scores are stable and associated with clinical outcome. (A)** Boxplots showing Olink module score comparisons of first available serum samples by Olink group. **(B)** Boxplots showing Olink module score comparisons of first available serum samples by final clinical outcome. **(C)** Heatmap showing discrete time Markov chain analysis probability of transition between Olink clusters and clinical outcome. For box plots, each dot represents a patient sample; center line, median; box limits, 25th and 75th percentile; whiskers, 1.5x IQR. Statistical significance (**A-B**) determined by 2-way ANOVA with Tukey's Multiple Comparisons correction. Adjusted p-values shown.

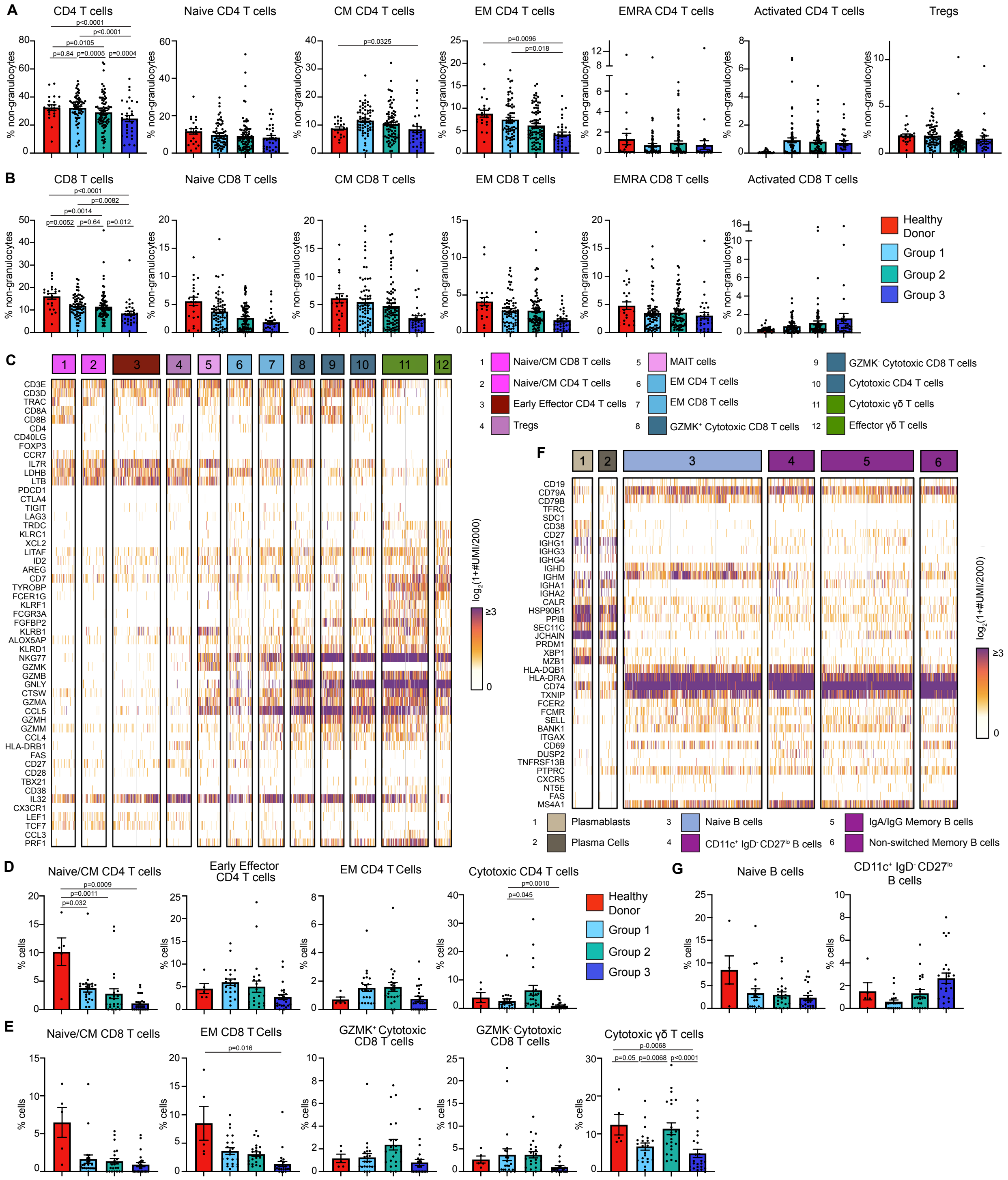

**Supplementary Figure 5. High dimensional characterization of circulating lymphocytes in COVID-19.** CD4 **(A)** and CD8 **(B)** T cell population frequencies (% non-granulocytes) in whole blood by Olink group measured by CyTOF. **(C)** Heatmap showing UMI counts of selected genes from T cell scRNAseq clusters from PBMC. Cell frequencies shown as % cells for select CD4 **(D)**, CD8 and γδ T cell **(E)** scRNAseq clusters. **(F)** Heatmap showing unique molecular identifier (UMI) counts of selected genes from B cell scRNAseq clusters from PBMC. **(G)** scRNAseq cell frequencies shown as % cells for select B cell clusters. For bar graphs, each dot (**A-B, D-E, G**) represents a patient sample. Statistical significance (**A-B, D-E, G**) determined by 2-way ANOVA with Holm-Sidak multiple comparisons correction. Adjusted p-values shown.

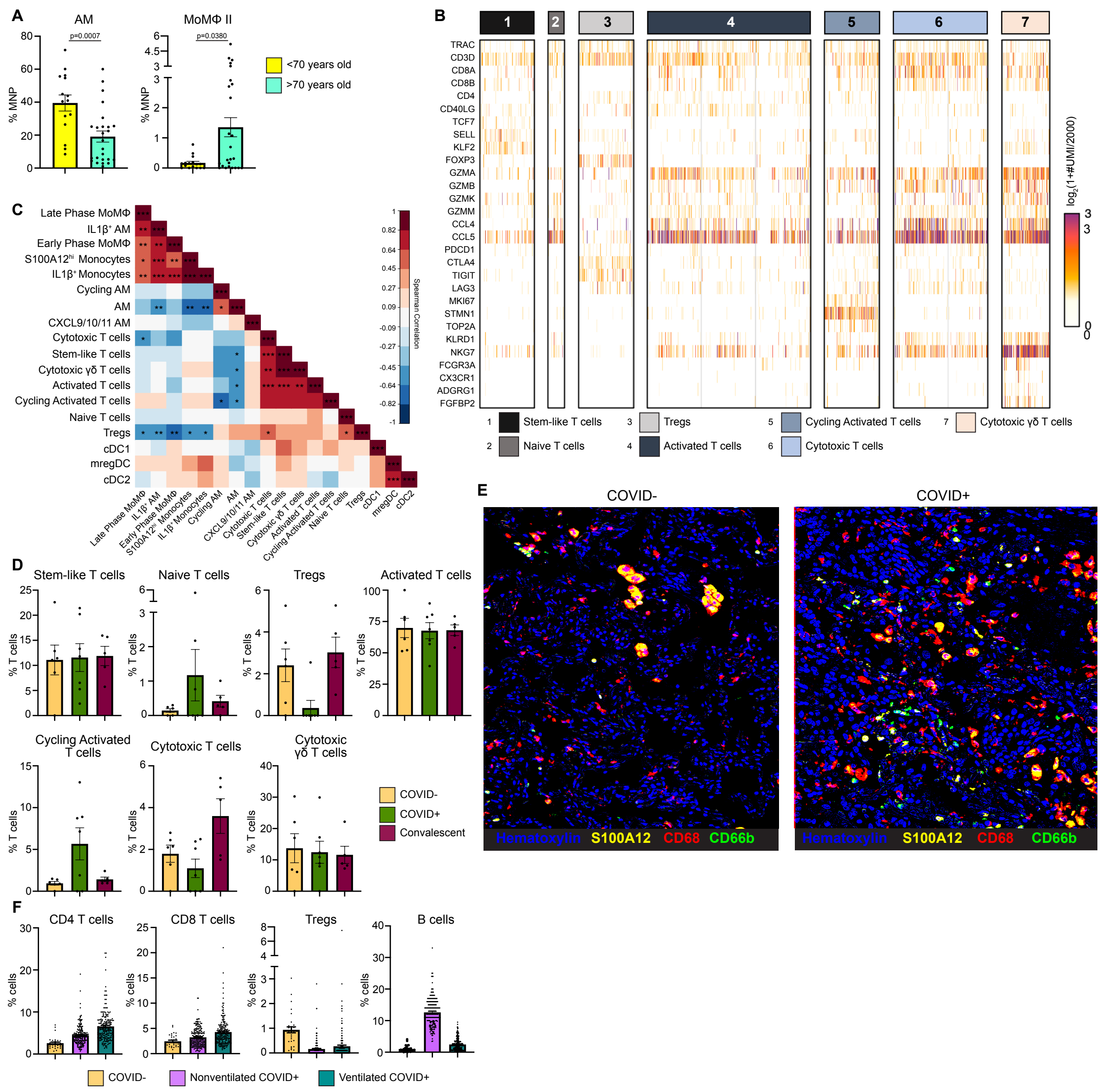

**Supplementary Figure 6. High dimensional characterization of immune cells in BAL (A)** AM and MoMΦ II frequencies as % MNP from scRNAseq analysis of normal lung tissue from untreated early stage non small cell lung cancer patients. Each dot represents single patient sample. Statistical significance determined by Mann-Whitney test. **(B)** Heatmap showing UMI counts of selected genes from T cell scRNAseq clusters from BAL. **(C)** Matrix of spearman correlation coefficients between identified scRNAseq BAL clusters. *p<0.05, **p<0.005, ***p<0.0005. **(D)** scRNAseq T cell cluster frequencies as % T cells from BAL. Each dot represents a patient sample **(E)** Overlaid, pseudocolored MICSSS image of COVID$^{+}$ and COVID$^{-}$ lungs, staining for S100A12, CD68, CD66b, and Hematoxylin. Granulocyte-like cells defined as CD66b$^{+}$ cells. **(F)** Quantification of lymphocytes in MICSSS images, shown as % cells. CD4 T cells defined as CD3$^{+}$ CD8$^{-}$ cells, CD8 T cells defined as CD3$^{+}$ CD8$^{+}$ cells, Tregs defined as CD3$^{+,}$ CD8$^{-}$ Foxp3$^{+}$ cells, B cells defined as CD20$^{+}$ cells.

**Supplementary Table 1. Mount Sinai COVID-19 Biobank Cohort.** IQR, Interquartile Range; EOD, end organ damage; CyTOF, cytometry by time of flight; scRNAseq, single cell RNA sequencing

| | **COVID+** | **COVID-** |
|---|---|---|
| Number of Subjects | 583 | 45 |
| Age (Median (IQR)) | 63 (53-74) | 36 (26-47) |
| Male Sex (%) | 61.2% | 46.7% |
| Deceased | 17.7% | 0.0% |
| **COVID-19 clinical severity at time of first sample** | | |
| Moderate COVID-19 | 61.1% | |
| Severe COVID-19 | 22.8% | |
| Severe COVID-19 with EOD | 12.3% | |
| N/A | 3.8% | |

**Supplementary Table 2. Mount Sinai Hospital Disease Severity Classification.** $SpO_2$, oxygen saturation; CXR, chest X-ray; CrCl, Creatinine Clearance; ALT, Alanine aminotransferase; ULN, upper limit of normal; RRT, renal replacement therapy

| **Disease Severity** | |
|---|---|
| Moderate COVID-19 | $SpO_2$<94% on RA or pneumonia by CXR, ≦6 L/min O2 support |
| Severe COVID-19 | >6L/min $O_2$ support, (-) pressors, CrCL>30 mL/min, ALT<5x ULN |
| Severe COVID-19 with EOD | >6 L/min $O_2$ support, (+) pressors, CrCl<30 mL/min or new RRT or ALT>5x ULN |

**Supplementary Table 3. Clinical information for PBMC scRNAseq samples.** PSO, post symptom onset; HD, healthy donor; EOD, Severe COVID-19 with End Organ Damage; CKD, chronic kidney disease

| Subject ID | Timepoint | Age | Sex | Days PSO | Disease severity | Olink Group | Olink Cluster | Final Clinical Outcome |
|---|---|---|---|---|---|---|---|---|
| PICR8016 | T1 | 28 | M | n/a | HD | n/a | n/a | Survived |
| PICR8017 | T1 | 25 | F | n/a | HD | n/a | n/a | Survived |
| PICR8106 | T1 | 58 | M | n/a | HD | 1 | 12 | Survived |
| PICR8107 | T1 | 56 | F | n/a | HD | 1 | 12 | Survived |
| PICR8108 | T1 | 50 | F | n/a | HD | n/a | n/a | Survived |
| PICR8109 | T1 | 51 | M | n/a | HD | n/a | n/a | Survived |
| PICR7250 | T1 | 68 | | n/a | n/a | 1 | 15 | Deceased |
| PICR7161 | T1 | 65 | M | 7 | EOD | 2 | 9 | Survived |
| | T12 | | | 19 | Moderate | 1 | 13 | |
| PICR7247 | T1 | 65 | F | 3 | Moderate | 1 | 13 | Survived |
| | T4 | | | 8 | Moderate | 1 | 14 | |
| PICR7114 | T1 | 80 | M | 32 | EOD | 2 | 6 | Survived |

| | | | | | | | | |
|---|---|---|---|---|---|---|---|---|
| | T4 | | | 35 | Moderate | 1 | 13 | |
| | T8 | | | 39 | Moderate | 1 | 15 | |
| PICR7393 | T1 | 83 | F | 16 | Severe | 1 | 13 | Survived |
| | T4 | | | 19 | Moderate | 2 | 6 | |
| PICR7061 | T12 | 89 | M | 19 | Severe | 1 | 13 | Survived |
| PICR7389 | T1 | 39 | F | 28 | Moderate | 1 | 14 | Survived |
| PICR7321 | T1 | 59 | M | 12 | Moderate | 1 | 14 | Survived |
| | T4 | | | 15 | Moderate | 2 | 7 | |
| | T8 | | | 21 | Moderate | 1 | 14 | |
| PICR7481 | T13 | 68 | M | 37 | Moderate | 1 | 14 | Survived |
| PICR7189 | T4 | 39 | M | 16 | Severe | 1 | 15 | Survived |
| | T8 | | | 20 | Severe | 1 | 15 | |
| PICR7162 | T4 | 47 | F | 11 | Severe | 1 | 15 | Survived |
| | T8 | | | 16 | Severe | 1 | 15 | |
| PICR7292 | T4 | 53 | M | 8 | Moderate | 1 | 15 | Survived |
| PICR7059 | T1 | 66 | M | 5 | Moderate | 2 | 8 | Survived |
| | T4 | | | 8 | Severe | 2 | 8 | |
| | T12 | | | 17 | Moderate | 2 | 8 | |
| | T13 | | | 27 | Moderate | 1 | 15 | |
| PICR7068 | T4 | 69 | F | 20 | Moderate | 1 | 15 | Survived |
| PICR7276 | T12 | 71 | M | 17 | Severe | 1 | 15 | Survived |
| PICR7066 | T1 | 70 | M | 11 | Moderate | 2 | 8 | Survived |
| | T4 | | | 17 | n/a | n/a | n/a | |
| PICR7420 | T1 | 78 | M | 29 | Moderate | 2 | 6 | Deceased |
| PICR7146 | T1 | 50 | F | 17 | Severe | 2 | 8 | Deceased |
| PICR7218 | T1 | 89 | F | 18 | Moderate | 3 | 5 | Survived |
| | T4 | | | 21 | Moderate | 2 | 6 | |
| PICR7403 | T1 | 20 | M | 5 | EOD | 3 | 3 | Survived |
| | T4 | | | 9 | Moderate | 2 | 7 | |
| | T8 | | | 12 | Moderate | 2 | 7 | |
| PICR7070 | T1 | 70 | M | n/a | Moderate | 2 | 7 | Survived |
| | T4 | | | n/a | Moderate | 2 | 7 | |
| PICR7484 | T1 | 66 | M | 9 | Severe | 2 | 8 | Survived |
| | T12 | | | 21 | EOD | 3 | 5 | |
| PICR7063 | T1 | 69 | M | 16 | Severe | 2 | 8 | Survived |
| | T8 | | | 23 | EOD | 3 | 5 | |
| PICR7333 | T1 | 32 | M->F | 7 | Severe | 2 | 9 | Survived |
| | T4 | | | 13 | Severe | 3 | 1 | |
| PICR7158 | T1 | 69 | F | 7 | Moderate | 2 | 9 | Survived |
| | T4 | | | 10 | EOD | 3 | 4 | |
| | T12 | | | 20 | EOD | 3 | 4 | |
| | T13 | | | 29 | EOD | 2 | 9 | |
| PICR7294 | T8 | 80 | M | 21 | Moderate | 2 | 9 | Survived |
| PICR7073 | T1 | 47 | M | 23 | EOD | CKD | 10 | Deceased |
| | T4 | | | 26 | EOD | CKD | 11 | |
| | T8 | | | 30 | EOD | 3 | 2 | |
| | T12 | | | 37 | EOD | 3 | 2 | |
| PICR7363 | T8 | 74 | F | n/a | EOD | 3 | 5 | Deceased |
| PICR7125 | T1 | 94 | F | 10 | EOD | 3 | 2 | Deceased |

| | | | | | | | | |
|---|---|---|---|---|---|---|---|---|
| PICR7239 | T1 | 68 | F | 14 | Severe | 3 | 3 | Deceased |
| PICR7405 | T4 | 84 | F | n/a | EOD | 3 | 5 | Deceased |
| PICR7292 | T1 | 53 | M | 5 | Moderate | 3 | 1 | Survived |
| PICR7091 | T4 | 66 | F | 9 | EOD | 3 | 2 | Survived |
| | T8 | | | 12 | Moderate | 3 | 2 | |
| | T12 | | | 16 | Moderate | 3 | 4 | |
| PICR7476 | T1 | 70 | F | 3 | EOD | 3 | 4 | Survived |
| | T4 | | | 6 | Moderate | 3 | 2 | |
| PICR7173 | T1 | 65 | M | 24 | EOD | 3 | 3 | Survived |
| | T4 | | | 27 | EOD | 3 | 3 | |
| PICR7175 | T4 | 69 | F | 6 | EOD | 3 | 4 | Survived |
| | T8 | | | 11 | EOD | 3 | 4 | |
| | T12 | | | 21 | Severe | 3 | 3 | |
| | T13 | | | 25 | Severe | CKD | 11 | |
| PICR7224 | T4 | 59 | F | 16 | Moderate | CKD | 10 | Deceased |
| | T8 | | | 20 | Moderate | CKD | 10 | |
| | T12 | | | 23 | Moderate | CKD | 10 | |
| PICR7178 | T1 | 33 | F | 9 | Moderate | CKD | 10 | Survived |
| PICR7266 | T4 | 57 | F | 36 | Moderate | CKD | 11 | Survived |
| | T8 | | | 40 | Moderate | CKD | 11 | |

**Supplementary Table 4. Clinical information for BAL samples.** PSO, post symptom onset; PI, post intubation

| Subject ID | COVID Status | Outcome | Days PSO | Days PI | Age | Sex |
|---|---|---|---|---|---|---|
| PICR7406 | + | Deceased | 10 | 2 | 77 | F |
| PICR7386 | + | Survived | 17 | 2 | 56 | M |
| PICR7295 | + | Deceased | 46 | 15 | 51 | M |
| PICR7022 | + | Survived | 41 | 2 | 51 | M |
| BAL8010 | + | Deceased | 22 | 1 | 72 | M |
| BAL8014 | + | Deceased | 18 | 0 | 94 | F |
| BAL8020 | + | Survived | 21 | 3 | 51 | F |
| BAL8007 | - | Survived | | | 61 | F |
| BAL8008 | Convalescent | Survived | | | 44 | M |
| BAL8009 | Convalescent | Survived | | | 51 | M |
| BAL8011 | - | Survived | | | 70 | F |
| BAL8012 | - | Survived | | | 65 | M |
| BAL8016 | - | Survived | | | 61 | M |
| BAL8017 | - | Survived | | | 53 | M |
| BAL8021 | Convalescent | Survived | | | 72 | M |
| BAL8022 | - | Survived | | | 59 | M |
| BAL8024 | Convalescent | Survived | | | 61 | M |
| BAL8027 | Convalescent | Survived | | | 62 | F |

**Supplementary Table 5. Clinical Information for lung autopsy samples.** PMI, post mortem interval; PSO, post symptom onset; AHRF, acute hypoxic respiratory failure; 2/2, secondary to

| Subject ID | Age | Sex | PMI (hours) | Days PSO | Days hospitalized | Intubated | Cause of Death |
|---|---|---|---|---|---|---|---|
| MA-20-120 | 59 | F | 5.5 | 13 | 8 | No | AHRF 2/2 COVID-19 |
| MA-20-149 | 77 | F | 7.5 | 25 | 22 | Yes | AHRF 2/2 COVID-19 |
| MA-20-123 | 61 | M | 3 | 29 | 48 | Yes | AHRF 2/2 COVID-19 |
| MA-20-81 | 57 | F | N/A | 4 | 4 | No | AHRF 2/2 COVID-19 |
